# Piezo2 voltage-block regulates mechanical pain sensitivity

**DOI:** 10.1101/2022.10.04.510762

**Authors:** Oscar Sánchez Carranza, Sampurna Chakrabarti, Johannes Kühnemund, Fred Schwaller, Valérie Bégay, Gary R. Lewin

**Author notes:** These authors contributed equally (Sampurna Chakrabarti, Johannes Kühnemund).

## Abstract

PIEZO2 mechanosensitive channels are required for normal touch sensation. However, PIEZO2 channels are almost completely blocked at negative resting membrane potentials. We show that PIEZO2 voltage-block can be relieved by mutations at a conserved Arginine (R2756) which dramatically sensitizes the channel to mechanical stimuli. We generated *Piezo2*^*R2756H/R2756H*^ and *Piezo2*^*R2756K/R2756K*^ *knock-in* mice to ask how voltage regulates the endogenous mechanosensitivity of sensory neurons. Surprisingly, mechanosensitive currents in nociceptors, neurons that detect noxious mechanical stimuli, were substantially sensitized in *Piezo2 knock-in* mice, but touch receptors were largely unaffected. *Piezo2 knock-in* mice were hypersensitive to noxious mechanical stimuli as their nociceptors acquired properties similar to ultrasensitive touch receptors. Thus, mechanical pain sensitivity can be tuned by voltage-block of PIEZO2 channels, a channel property potentially amenable for pharmacological modulation.

## Introduction

*Piezo2* is genetically required for normal touch sensation (Chesler et al., 2016; Ranade et al., 2014; Coste et al., 2010, 2012), and it is widely assumed that PIEZO2 channels form the conduction pore of native mechanosensitive currents that underlie touch receptor mechanosensitivity. However, despite the fact that *Piezo2* channels are expressed in almost all sensory neurons of the dorsal root ganglia (DRG), deletion of *Piezo2* leads to complete loss of mechanosensitivity only in around half of all mechanoreceptors (Ranade et al., 2014; Murthy et al., 2018). Additionally, the mechanosensitivity of almost all nociceptors is preserved in the absence of *Piezo2* (Murthy et al., 2018). Work in nematodes has shown how genetic deletion of candidate mechanotransduction channels does not always provide definitive evidence that the protein forms the pore of the native mechanosensitive current (Geffeney and Goodman, 2012; O’Hagan et al., 2005; Goodman and Sengupta, 2019). A powerful way to directly assess the participation of a channel in transduction is to change the biophysical properties of the endogenous channel with the prediction that native mechanosensitive currents should acquire these new biophysical properties (O’Hagan et al., 2005). PIEZO channels are not only gated by mechanical stimuli, but are also controlled by membrane voltage. Thus, at physiological membrane potentials >90% of PIEZO channels cannot be opened by mechanical stimuli, but are made available at depolarized membrane potentials (Moroni et al., 2018). We previously identified a single highly conserved Arginine residue in PIEZO1 channels (mR2482) that when mutated effectively eliminates most of the PIEZO1 voltage-block (Moroni et al., 2018) (Fig. 1a). Interestingly, mutations in the same conserved residue of PIEZO2 (mPIEZO2; R2756; hPIEZO2 R2686) are associated with distal arthrogryposis, Gordon syndrome and the Marden-Walker Syndrome, all of which are human developmental disorders (Alisch et al., 2016; Mcmillin et al., 2014). Here we show a single mutation can abrogate the voltage-block of the PIEZO2 channel, dramatically increasing channel availability at physiological membrane potentials. By mutating the same site in the channel in vivo we could investigate the effects of changing the channel properties on native mechanosensitive currents and their effects on sensory physiology. Surprisingly, we observed only minor effects on touch receptors, but the properties of mechanosensitive currents in nociceptors were dramatically sensitized in a way that reflected the changes in PIEZO2 channel function. Our data show how the voltage block of PIEZO2 serves to keep the mechanical threshold of nociceptors high so that they detect noxious and not non-noxious mechanical stimuli. Furthermore, our data suggest a simple model whereby different kinds of algogens drive nociceptor sensitization by releasing PIEZO2 channels from voltage-block.

**Figure 1.**
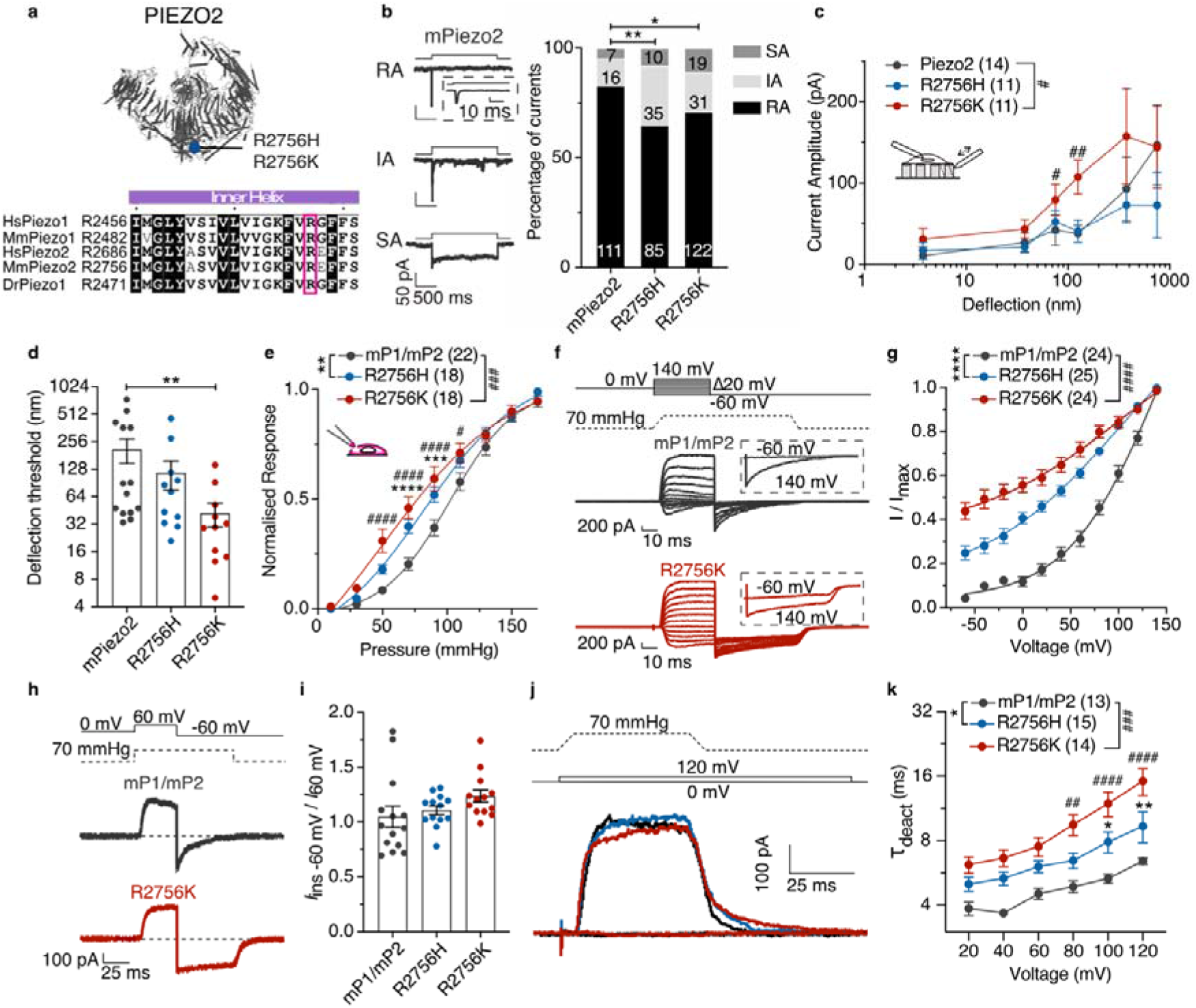
Mutations in the R2756 of *mPiezo2* are gain of function mutations. (**a**) *Above*, structural model of PIEZO2 (PBD ID: 6KG7(Wang et al., 2019)) indicating the position of the R2756 (blue dot). *Below*, residues alignment (using ESPript 3.0(Robert and Gouet, 2014)) showing that the Arginine (pink square) is conserved in PIEZO channels. (**b**) *Left*, Example traces of rapidly adapting (RA), intermediate adapting (IA) and slowly adapting (SA) currents from N2a^Piezo1-/-^ cells overexpressing mPiezo2. *Right*, Proportion of RA currents is decreased in cells expressing mPiezo2 variants. Numbers represent the currents recorded (χ^2^ test, *P=0.04, **P=0.004). (**c**) Deflection-response relationships showing that R2756K mutant is more sensitive to mechanical stimuli compared to wild type (Mann Whitney test ^#^P=0.04, ^##^P=0.008; Two-way ANOVA indicated differences between variants, ^#^P=0.04). (**d**) Deflection threshold was lower in the R2756K mutant (Kruskal-Wallis test, **P=0.006). (**e**) Stretch-response curves of the chimeric channel variants. Peak currents were normalised according to their maximum (Two-way ANOVA **P= 0.003, ^###^P=0.0002, with Sídák post hoc test ***P=0.0006, ****P<0.0001, ^#^P=0.03, ^####^P<0.0001). (**f**) Representative traces of the tail current protocol performed in N2a^Piezo1-/-^ cells expressing the chimeric channels. *Insert*, tail currents evoked after pre-stimuli of -60 and 140 mV. (**g**) The apparent open probability increased in the mutants at physiological pulses. Tail currents were normalised to their maximum (Two-Way ANOVA, Sídák test, ****P<0.0001, ^####^P<0.0001). (**h**) Example traces of the rectification index (*I*_ins -60 mV_ / *I*_60 mV_) protocol. (**i**) Rectification index is similar in the mutants. **(j**) Representative traces from the deactivation protocol in the chimeric channel variants. (**k**) Mutants displayed slower deactivation kinetics at depolarizing pulses. An exponential fit was calculated to measure the deactivation time (τ_deact_; Two-Way ANOVA, Sídák test, *P<0.05, **P=0.007, ^##^P=0.002, ^###^P=0.0002, ^####^P<0.0001). Data are presented as mean ± s.e.m.

## Results

We first asked whether the conserved R2756 residue also controls the voltage sensitivity of mPIEZO2 channels. We thus generated *mPiezo2* channels with single missense mutations (R2756H, R2756C and R2756K), known to be associated with human developmental diseases. We quantitatively assessed mechanosensitivity using substrate deflection of N2a^Piezo1-/-^ cells expressing wild type or mutant *Piezo2* channels (Moroni et al., 2018; Poole et al., 2014; Servin-Vences et al., 2017) (Supplementary Fig. 1a-c). We measured three types of mechanically gated currents in cells expressing *Piezo2* channels: rapidly adapting (RA), intermediate adapting (IA) and slowly adapting currents (SA) (Fig. 1b, Supplementary Fig. 1d). Cells expressing the R2756H, R2756C and R2756K mutations exhibited significantly fewer RA and increased proportions of IA and SA currents compared to wild type (Fig. 1b, Supplementary Fig. 1d). The deflection-current relationship revealed that R2756K mutant channels are more sensitive to pilli deflection compared to wild type or R2756H/R2756C mutant channels (Fig. 1c, Supplementary Fig. 1e). Consistently the mean deflection threshold for R2756K was almost five-fold lower than that of wild type or R2756H/R2756C mutant channels (Fig. 1d, Supplementary Fig. 1f). We also noted subtle, but significant changes in the kinetics of mechanosensitive currents generated by *Piezo2* mutant channels (e.g. small increase in latency for activation) (Extended data Table 1). We next measured the effects of these mutations on the stretch and voltage sensitivity of PIEZO2 channels. Mutations were introduced into the stretch-sensitive chimeric channel mP1/mP2(Moroni et al., 2018) and currents were measured from excised outside-out patches. The R2756K and R2756H mutant channels displayed significantly enhanced stretch sensitivity compared to wild type chimeric channels, but the R2756C substitution did not alter stretch sensitivity (Fig. 1e, Supplementary Fig. 2a,b). Additionally, the R2756K and R2756H chimeric variants showed significantly slower inactivation kinetics compared to wild type channels (Supplementary Fig. 2d). We next used a tail current protocol to measure channel availability(Moroni et al., 2018) (Fig. 1f). Between 25 and 45% of the maximum tail current could be measured from the R2756H and R2756K chimeric variants at -60 mV compared to less than 5% in wild type (Fig. 1f-g, Supplementary Fig. 3a). Thus, both R2756H and R2756K mutations substantially relieve the voltage-block of these chimeric channels at physiological membrane potentials. The effect on the tail current was not accompanied by any change in the rectification index (Fig. 1h-i). PIEZO2 channels inactivate very rapidly at negative potentials making it challenging to study deactivation kinetics. We thus measured the effects of pressure removal at a series of positive voltages on current deactivation and found that R2756H and R2756K chimeric variants showed significantly slower deactivation compared to the wild type chimera (Fig. 1j-k). A considerable delay in channel closing was also observed during the transition from the inactivated to deactivated state after pressure removal (Supplementary Fig. 3b,d). In conclusion, slower inactivation and deactivation, increased mechanosensitivity and an almost complete removal of voltage-block were the main effects of the R2756H and R2756K *mPiezo2* missense mutations, with the R2756K mutation clearly displaying the strongest effects on all these parameters.

**Figure 2.**
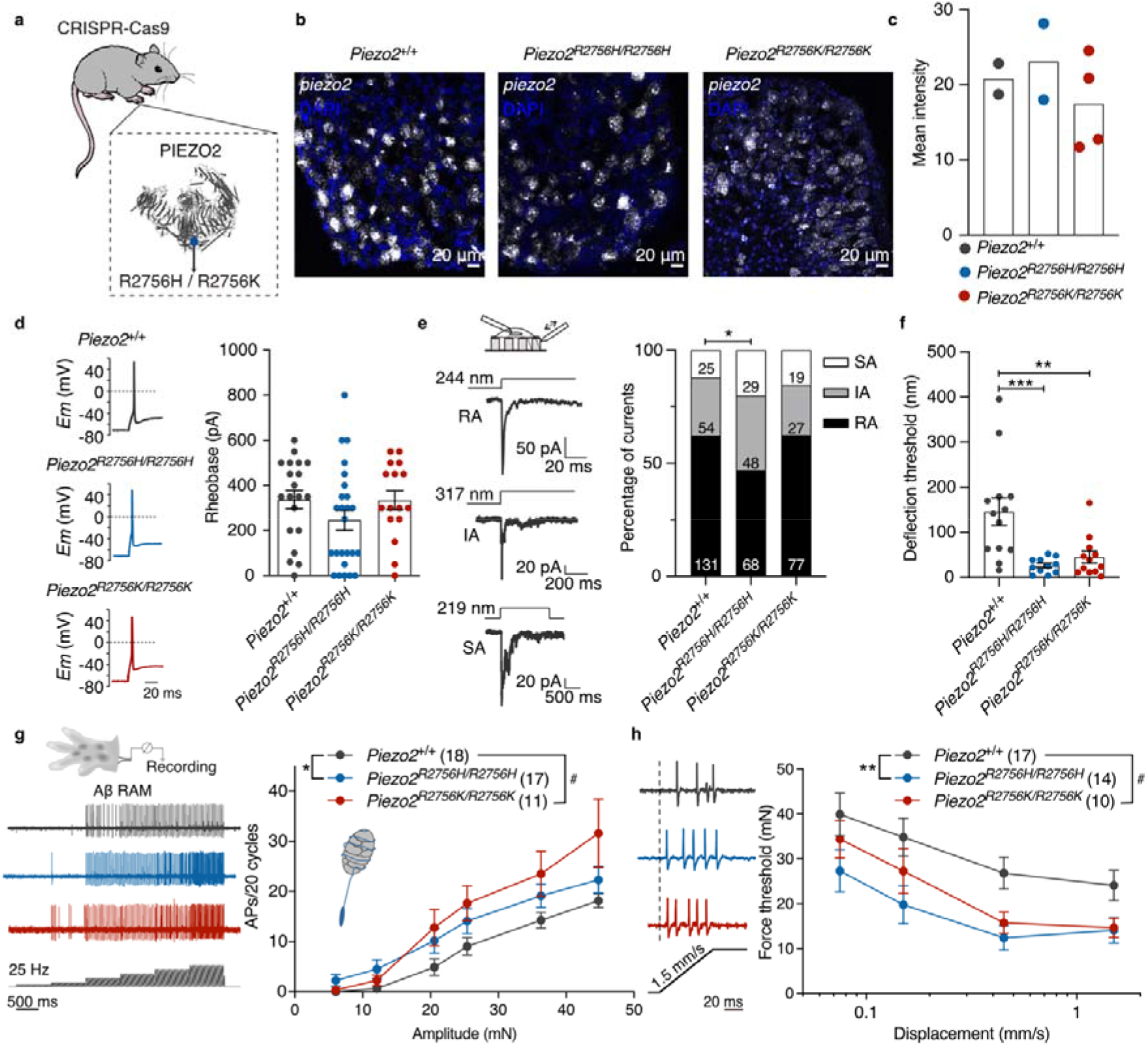
Mechanoreceptors from *knock-in* mice are more sensitive to deflection stimuli. (**a**) Cartoon representing the global insertion of the mutations in *Piezo2 knock-in* mice. (**b**) Representative images of *Piezo2 in-situ* hybridization (RNAscope) in lumbar DRGs. White, *Piezo2* mRNA; blue, 4’,6-diamidino-2-phenylidole (DAPI). (**c**) Quantification of area of transcript fluorescence from all sections. Each dot represents the mean value from each mouse. (**d**) *Left*, Representative APs in mechanoreceptors from *Piezo2*^+/+^, *Piezo2*^*R2756H/R2756H*^ and *Piezo2*^*R2756K/R2756K*^ animals. *Right*, Distribution of AP thresholds (Rheobase) showing no differences between mutants and wild type. (**e**) *Left*, Representative traces of the three types of deflection-gated currents (RA, IA and SA) from *Piezo2*^+/+^ mechanoreceptors. *Right*, Histograms showing that *Piezo2*^*R2756H/R2756H*^ neurons displayed less RA currents compared to wild type cells (χ^2^ test, *P=0.01). Numbers indicate the total of currents recorded. (**f**) Deflection thresholds were lower in *Piezo2*^*R2756H/R2756H*^ and *Piezo2*^*R2756K/R2756K*^ cells compared to wild type (Kruskal-Wallis test, **P=0.008, ***P=0.0008). (**g**) *Left*, Example traces of single RAM Aβ fibres form wild type and *Piezo2 knock-in* mice in response to a 25 Hz vibration stimulius. *Right*, Increased AP firing was observed in RAM Aβ fibres from the mutants compared to wild type (Two-way ANOVA with Sídák post hoc analysis, *P=0.04, ^#^P=0.01). (**h**) *Left*, Representative ramp responses of individual RAMs from *Piezo2 knock-in* and wild type mice at 1.5 mm/s. *Right*, Displacement-thresholds relationships showing that RAM Aβ fibres from Piezo2 mutants are more sensitive to mechanical stimuli (Two-way ANOVA with Sídák post hoc analysis, **P=0.007, ^#^P=0.03). Data are presented as mean ± s.e.m.

**Figure 3.**
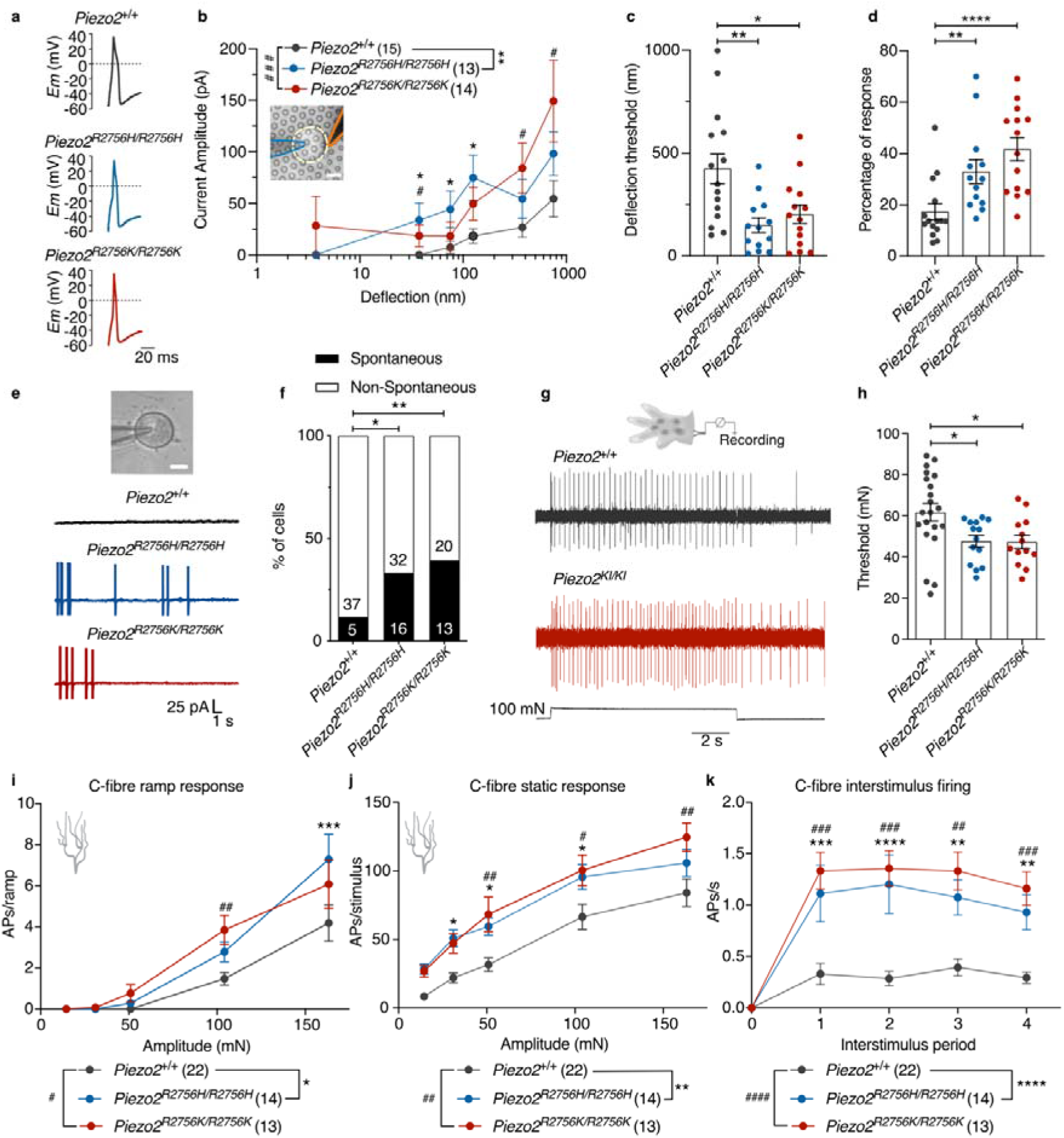
Nociceptors from knock-in mice showed spontaneous AP firing. (**a**) Representative APs in nociceptors from *Piezo2*^+/+^, *Piezo2*^*R2756H/R2756H*^ and *Piezo2*^*R2756K/R2756K*^ animals. (**b**) Deflection-current amplitude relationship of nociceptors showing that neurons from *knock-in* displayed hypersensitive deflection-gated currents. (Mann Whitney test *P<0.05, ^#^P<0.05; additionally, an ordinary Two-way ANOVA indicated differences between wild type and the mutants, **P=0.005; ^###^P=0.0008). (**c**) Deflection thresholds were lower in nociceptors from *knock-in* mice compared to wild type (one-way ANOVA, *P=0.01, **P=0.002). (**d**) Histogram showing that nociceptors from *Piezo2*^*R2756H/R2756H*^ and *Piezo2*^*R2756K/R2756K*^ are more responsive to deflection stimuli compared to controls (Kruskal-Wallis test, **P=0.005, ****P<0.0001). (**e**,**f**) Representative traces of current clamp recordings and percentage of neurons showing spontaneous activity in nociceptors from control and *knock-in* mice. Numbers indicate number of cells recorded (χ^2^ test, *P=0.01, **P=0.005). (**g**) *Top*, cartoon representing *ex-vivo* glabrous skin-nerve preparation. *Bottom*, Representative traces of C-fibre activity during a 100 mN stimulus from control (*Piezo2*^+/+^) and Piezo2 *knock-in* (*Piezo2*^*KI/KI*^) animals. (**h**) Histogram showing that C-fibres from mutants are more sensitive to mechanical stimuli compared to wild type (One-way ANOVA, *P<0.05). (**i-k**) Spike activity during ramp phase (**i**, Two-way ANOVA with Sídák post hoc analysis, *P=0.02, ^#^P=0.02, ^##^P=0.007, ***P<0.001), static phase (**j**, Two-way ANOVA with Sídák post hoc analysis, *P<0.05, ^#^P<0.05, **P=0.003, ^##^P<0.01) and interstimulus firing (**k**, Two-way ANOVA with Sídák post hoc analysis, **P<0.01, ^##^P<0.01, ^***^P<0.001, ^###^P<0.001, ^****^P<0.0001, ^####^P<0.0001). Data are presented as mean ± s.e.m.

Our biophysical measurements led us to predict that introduction of R2756H and R2756K into the mouse genome should radically alter the mechanosensitivity of endogenous PIEZO2-dependent currents. We generated two *knock-in* mice that globally express the R2756H and R2756K variants (*Piezo2*^*R2756H*^ and *Piezo2*^*R2756K*^ mice) (Fig. 2a, Supplementary Fig. 4a,b). The orthologous human mutation of *Piezo2*^*R2756H*^ has been associated with short stature and scoliosis (Alisch et al., 2016; Haliloglu et al., 2017; Mcmillin et al., 2014). Interestingly, we found that homozygous *Piezo2*^*R2756H/R2756H*^ animals weighed on average ∼20% less than wild type controls at 4 weeks of age. At 8 and 12 weeks of age, both *Piezo2*^*R2756H/R2756H*^ and *Piezo2*^*R2756K/R2756K*^ mice weighed significantly less on average than wild types (∼9% less), however, this effect was only partially penetrant as many of the mutant mice had body weights in the same range as controls. No effect of the mutations on body weight were observed in heterozygous animals (Supplementary Fig. 4c,d). In 50% of the *Piezo2*^*R2756K/R2756K*^ mice (9/18) we observed abnormal spine curvature (scoliosis), but this phenotype was not observed in heterozygotes or in *Piezo2*^*R2756H/R2756H*^ mutant mice (Supplementary Fig. 4e).

**Figure 4.**
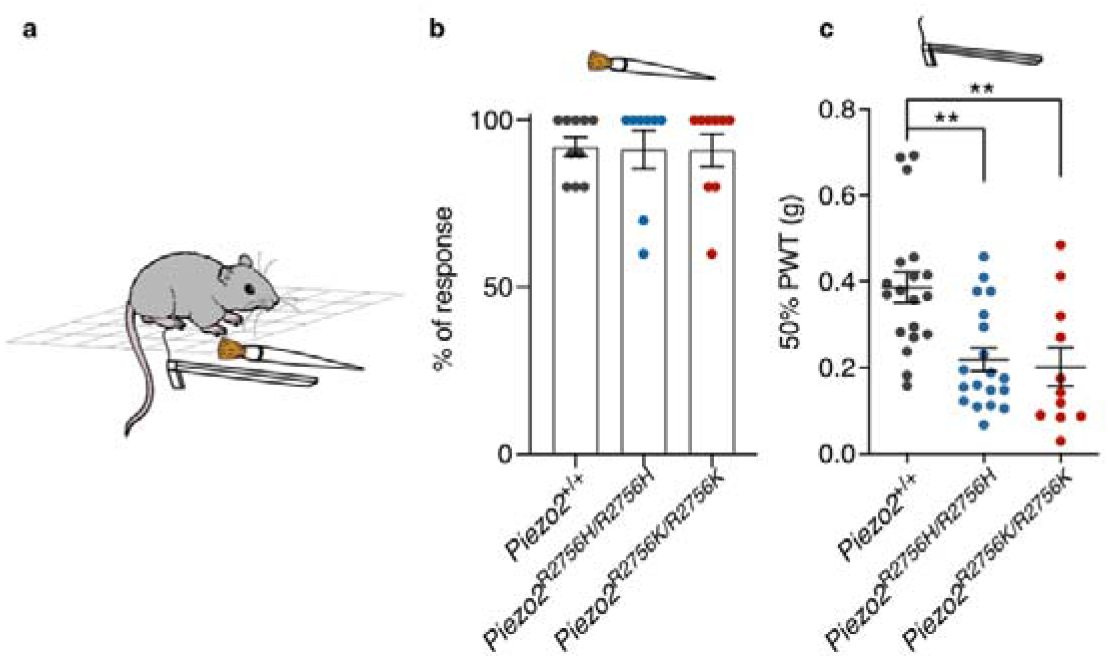
Mechanical pain hypersensitivity in *Piezo2* knock-in mice. (**a**) Cartoon representing the brush and von Frey experiments. When mice withdrew or licked their paw, the stimulation was considered as a positive response. (**b**) Histogram showing the percentage of response to brush stimulation in mutants (*Piezo2*^*R2756H/R2756H*^, n=8; *Piezo2*^*R2756K/R2756K*^, n=9) and wild type (n=10) (**c**) *Piezo2*^*R2756H/R2756H*^ (n=19) and *Piezo2*^*R2756K/R2756K*^ (n=11) animals showed a reduced 50% PWT compared to controls (n=19) (One-way ANOVA test, **P=0.001). Each dot represents average values from different measures taken on different days in each animal. Data are presented as mean ± s.e.m.

The introduction of missense mutations could alter gene expression, we thus examined *Piezo2* expression in sensory neurons within the dorsal root ganglia (DRG) using RNAscope. We found that in the DRG *Piezo2*^*+/+*^, *Piezo2*^*R2756H/R2756H*^ and *Piezo2*^*R2756K/R2756K*^ mice showed similar *Piezo2* mRNA levels (Fig. 2b, c). In the complete absence of *Piezo2*, around half of mechanoreceptors are completely insensitive to mechanical stimuli (Ranade et al., 2014; Murthy et al., 2018). We next recorded mechanosensitive currents in wild type and mutant sensory neurons in culture which had been classified as mechanoreceptors or nociceptors according to their size and action potential (AP) shape as previously reported (Lechner et al., 2009; Poole et al., 2014; Rose et al., 1986; Koerber et al., 1988) (Fig. 2d, Supplementary Fig. 3a). When performing current clamp recordings from mechanoreceptors we found their excitability to be unaffected by the *Piezo2* point mutations as reflected by unaltered resting membrane potentials, rheobase or input resistance (Fig. 2d, Supplementary Fig. 5a, Extended data Table 2). We next recorded deflection gated currents from mechanoreceptors and again identified mechanically activated currents with RA, IA and SA kinetics, with RA-currents predominating (Poole et al., 2014; Hu and Lewin, 2006). Mechanoreceptors from *Piezo2*^*R2756H/R2756H*^ and *Piezo2*^*+/R2756K*^ showed a small but significant decrease in the proportion of RA currents compared to wild type cells, but no significant differences were observed in mechanoreceptors from *Piezo2*^*R2756K/R2756K*^ and *Piezo2*^*+/R2756H*^ mice (Fig. 2e, Supplementary Fig. 6b). Deflection-current amplitude relationships were similar between genotypes with a trend for mechanoreceptors from *Piezo2*^*R2756K/R2756K*^ mice to show higher sensitivity (Supplementary Fig. 6a,c). However, we observed robust and statistically significant reductions in the mean minimum deflection amplitudes capable of evoking mechanosensitive currents in all homozygous and heterozygous variant genotypes compared to wild type (Fig. 2f, Supplementary Fig. 6d). This change in threshold was accompanied by small changes in the kinetic parameters of mechanosensitive currents, for example in the inactivation kinetics of RA-currents in *Piezo2*^*R2756H/R2756H*^ mutants (Extended data Table 2). We next asked if the threshold for gating the mechanosensitive currents in isolated sensory neurons was accompanied by changes in the properties of intact mechanoreceptors. Using an *ex-vivo* preparation we recorded single mechanoreceptors innervating the hind paw glabrous skin (Schwaller et al., 2021; Walcher et al., 2018a). We found that rapidly-adapting mechanoreceptors (RAMs) from *Piezo2*^*R2756H/R2756H*^ and *Piezo2*^*R2756K/R2756K*^ mutants that innervate Meissner’s corpuscles displayed mildly enhanced firing to small 25 Hz sinusoidal stimuli compared to wild type (Fig. 2G). During the ramp phase of the mechanical stimulus RAMs recorded from *Piezo2*^*R2756H/R2756H*^ and *Piezo2*^*R2756K/R2756K*^ fired with shorter latencies reflecting lower force thresholds that were up to 10 mN smaller compared to wild type mice (∼50% reduction) (Fig. 2h, Supplementary Fig. 7a,b). In contrast, slowly-adapting mechanoreceptors (SAMs) associated with Merkel cells (Woo et al., 2014; Maksimovic et al., 2014) were barely affected by either missense mutation (Supplementary Fig. 7c-f). Thus, a sub-population of mechanoreceptors had significantly altered receptor properties when the biophysical properties of PIEZO2 are altered. This data is consistent with the idea that other unknown mechanosensitive channels regulate the sensitivity of many mechanoreceptors.

We were surprised by the fact that large changes in the biophysical properties of endogenous PIEZO2 channels only had mild effects on touch receptors. However, there is increasing evidence that PIEZO2 may also play a role in pain sensitivity (Murthy et al., 2018; Szczot et al., 2018). Nociceptors that detect intense mechanical stimuli do not lose mechanosensitivity in the absence of PIEZO2, but show reduced initial firing to step mechanical stimuli (Murthy et al., 2018). *Piezo2* is expressed by most nociceptors and so we next examined the effects of *Piezo2* missense mutations on nociceptor physiology. We measured the mechanosensitivity of nociceptive sensory neurons with broad humped action potentials (Fig. 3a). We found that the deflection evoked currents were often three times larger at all deflection amplitudes in neurons from *Piezo2*^*R2756H/R2756H*^ and *Piezo2*^*R2756K/R2756K*^ mice compared to wild type cells (Fig. 3b). In addition, the threshold for current activation was substantially lowered to values typical of mechanoreceptors in both types of mutant neurons (Fig. 3c, Fig. 2f). The frequency with which a mechanical stimulus evoked currents was also substantially increased in mutant neurons compared to wild type (Fig. 3d). We also noted significant, but much milder, increases in the sensitivity of deflection evoked currents in neurons from animals in which either mutation was present on only one allele (Supplementary Fig. 8a-c). Normally, acutely cultured sensory neurons exhibit little or no ongoing action potential firing (Chakrabarti et al., 2020). Interestingly, we found that nociceptors from *Piezo2*^*R2756H/R2756H*^ and *Piezo2*^*R2756K/R2756K*^ often exhibited ongoing firing in the absence of current injection compared to wild type neurons (Fig. 3e,f). Moreover, we measured the rheobase of neurons from *Piezo2*^*R2756H/R2756H*^ and *Piezo2*^*R2756K/R2756K*^ animals and found this to be decreased by 30% and 55%, respectively compared to wild type (Supplementary Fig. 9a,b). Such changes in electrical excitability could be due to alterations in voltage-gated conductance, however direct measurements of macroscopic voltage-gated inward and outward currents revealed no significant differences between wild type and mutant neurons (Supplementary Fig. 9c,d). The resting membrane potential of mutant neurons was also not altered compared to wild type (Extended data Table 3). Thus, nociceptors from *Piezo2*^*R2756H/R2756H*^ and *Piezo2*^*R2756K/R2756K*^ mice exhibit substantial mechanical hyperexcitability. We next examined intact nociceptors innervating the skin of which there are two main classes, thinly myelinated Aδ-fibers or unmyelinated C-fibers nociceptors (Lewin and Moshourab, 2004). In the glabrous skin we observed using quantitative force stimuli that C-fiber nociceptors from *Piezo2*^*R2756H/R2756H*^ and *Piezo2*^*R2756K/R2756K*^ mice exhibited substantially increased firing rates and much lower mechanical thresholds for activation compared to wild type (Fig. 3g-j) as did Aδ-fiber mechanonociceptors (Supplementary Fig. 10a-d). We analysed firing during stimulus onset (ramp phase) separately from the static phase and found that there was a substantial sensitization to both phases in C-fiber and Aδ-fiber mechanonociceptors in both mutant mice (Fig. 3i,j Supplementary Fig. 10 and 11). A hallmark feature of C-fiber nociceptors, first described by Perl in the 1960s, is that they often continue to fire after the noxious mechanical stimulus is removed (Bessou and Perl, 1969). Strikingly, C-fibers recorded from the mutants showed substantially increased ongoing firing after removal of mechanical stimuli compared to wild type C-fibers (Fig. 3g). We quantified this change in C-fibers from *Piezo2*^*R2756H/R2756H*^, *Piezo2*^*+/R2756K*^, *and Piezo2*^*R2756K/R2756K*^ mice and found that in these genotypes C-fibers exhibited up to three-fold increased interstimulus firing activity compared to wild type (Fig. 3k, Supplementary Fig. 11c). Only in *Piezo2*^*+/R2756H*^ mice, was the interstimulus firing was equivalent to that seen in wild type controls. These *in-vitro* and *ex-vivo* data show that voltage control of PIEZO2 channels in nociceptors is crucial for conferring high mechanical thresholds to mammalian nociceptors. Furthermore, the impairment in the ability of mutant channels to deactivate after opening was correlated with a large increase in ongoing activity of nociceptors in the absence of a mechanical stimulus.

Apart from a non-penetrant scoliosis or occasional growth retardation, especially in *Piezo2*^*R2756K/R2756K*^ mice, the *Piezo2 knock-in* mice appeared largely healthy, with no obvious motor deficits. We tested behavioral responses to innocuous brushing of the hindpaw and found no obvious hypersensitivity in *Piezo2*^*R2756H/R2756H*^ and *Piezo2*^*R2756K/R2756K*^ mice compared to wild type controls (Fig. 4a). However, paw withdrawal to punctate stimulation were clearly sensitized with paw withdrawal thresholds (PWT)(Chaplan et al., 1994; Christensen et al., 2020; Dixon, 1980) on average half those of controls in both *Piezo2*^*R2756H/R2756H*^ and *Piezo2*^*R2756K/R2756K*^ mutant genotypes (Fig 4c). Thus, relief of the voltage-block of Piezo2 is associated with heightened mechanical pain sensitivity *in-vivo*.

## Discussion

Here we have shown that the biophysical properties of PIEZO2 channels sets the sensitivity and mechanical thresholds of nociceptors required to detect painful mechanical stimuli. Changing PIEZO2 residue R2756 to histidine or lysine made nociceptors approximately 3-fold more sensitive to mechanical stimuli with mechanical thresholds similar to touch receptors. This remarkable change in excitability resembled physiological sensitization processes that follow strong chemical or mechanical activation of nociceptors in humans, primates and rodents (Lewin and Moshourab, 2004; Kress et al., 1992; Schmidt et al., 1995; Meyer et al., 1991; Steen et al., 1995). The observed changes in mechanosensitive currents measured from recombinant ion channels or isolated nociceptors were remarkably predictive of changes in the *in-vivo* sensitivity of nociceptors. For example, R2756 mutations dramatically slowed the closing of PIEZO2 channels a phenomenon that was reflected in ongoing activity of nociceptors after the cessation of the mechanical stimulus. We also show that R2756 mutations strongly influence the excitability of C-fiber nociceptors so that spontaneous activity was seen both *in-vitro* and *in-vivo*. This was a very surprising finding as it shows for the first time that it is not only voltage gated sodium channels like Na_V_1.7, Na_V_1.8 or Na_V_1.9 that have the potential to control nociceptor excitability (Bennett et al., 2019), but also mechanosensitive channels that are controlled by membrane voltage. Activation of nociceptors by inflammatory mediators or algogens, like capsaicin (Murthy et al., 2018), will strongly depolarize sensory endings thus potentially relieving voltage block of PIEZO2 channels. Membrane depolarization, which in the very compact nociceptor ending may be considerable, has thus the potential to mimic relief of voltage block that keeps the threshold to activate nociceptors high. Thus, we propose that voltage control of abundant PIEZO2 channels in most nociceptors is a major final mediator of nociceptor sensitization caused by strong nociceptive stimuli. In contrast to nociceptors, changing the biophysical properties of PIEZO2 was associated with only minor changes in the threshold and suprathreshold sensitivity of some, but not all touch receptors. The contrast between nociceptors and mechanoreceptors was striking and further supports the idea that uncharacterized mechanosensitive channels underlying touch sensation remain to be identified.

## Methods

All experiments with mice were done in accordance with protocols reviewed and approved by the German Federal authorities (State of Berlin).

### Molecular Biology

DNA constructs containing mPiezo2, mP1/mP2 and the variants were purified from transformed bacteria grown in large-scale bacterial culture (50 mL Midiprep, PureYield™ Plasmid Midiprep System, Promega). The midipreps were made according to the manufacture’s protocol. DNA quantification was measured using a NanoDrop 2000 (Thermofisher Scientific).

Insertion of point mutations in mPiezo2 and the chimeric channel mP1/mP2 were carried out using the Q5® Site-Directed Mutagenesis Kit (NEB, Inc) according to the manufacture’s indications. Specific primers for each mutant were used at 0.5 µM. Variant R2746H was generated using forward primer 5’-GAAATTTGTTCATGAGTTCTTCAG-3’, R2756C using forward primer 5’-GAAATTTGTTTGTGAGTTCTTCAG-3’ and R2756K using forward primer 5’-GAAATTTGTTAAAGAGTTCTTCAGTGGG-3’. For all mutants the same reverse primer was used, 5’-CCAATTACAAGGACAACAG-3’. Polymerase chain reactions (PCR) products were used as template for bacteria transformation and ampicillin resistant colonies were chosen and grown in large-scale bacterial culture for DNA purification. DNA plasmids were sequenced to verify the insertion of point mutations.

### Cultured cell lines

N2a^Piezo1-/-^ cells(Moroni et al., 2018) were used for the characterisation of the biophysical properties of the mutants of mPiezo2 and the chimeric channel mP1/mP2. Recently we showed that these cells lack stretch-and deflection-gated currents(Moroni et al., 2018; Patkunarajah et al., 2020). Cells were cultured in 45% DMEM-Glutamax (gibco, ThermoFisher SCIENTIFIC), 45% Opti-MEM (gibco, ThermoFisher SCIENTIFIC), 10% fetal calf serum (PAN Biotech GMBH) and 1% penicillin and streptomycin (Sigma-Aldrich) media.

N2a^Piezo1-/-^ cells were transiently transfected using FuGeneHD (Promega, Madison). A mix of 100 μL of Opti-MEM, 3 μL of FuGeneHD and 1 μg of DNA was incubated for 10 min at room temperature. The mix was added to Na2^Piezo1-/-^ cells cultured in 30 mm x 15 mm Petri dishes and 900 μL of media containing 50% DMEM-Glutamax and 50% Opti-MEM was added for an overnight transfection. Electrophysiological recordings were made 18-24 h post-transfection. At least four transfections were made for each set of experiments.

### DRG culture

DRG neurons were collected from all the spinal segments in plating medium on ice (DMEM-F12 (Invitrogen) supplemented with L-Glutamine (2 μM, Sigma-Aldrich), Glucose (8 mg/ml, Sigma Aldrich), Penicillin (200 U/mL)-Streptomycin (200μg/mL) and 10 % fetal horse serum). The DRGs were treated with Collagenase IV (1 mg/ml, Sigma-Aldrich) for 1 h at 37°C and then washed three times with Ca^2+^-and Mg^2+^-free PBS. The samples were incubated with trypsin (0.05%, Invitrogen, Karlsruhe) for 15 min, at 37°C. After the enzymatic treatment, the collected tissue was triturated with a pipette tip and plated in a droplet of plating medium on the elastomeric pillar arrays precoated with laminin (4 μg/cm^2^, Invitrogen) as described in Poole K., et al. for the pillar arrays experiments (see preparation of pillar arrays section)(Poole et al., 2014). Cells were cultured overnight, and the electrophysiology experiments were preformed after 18-24 h of the dissection.

### Preparation of pillar arrays

Pillar arrays were prepared as previously described(Poole et al., 2014; Servin-Vences et al., 2017; Patkunarajah et al., 2020). Briefly, silanized negative masters were used as templates. Negative masters were covered with polydimethylsilozane (PDMS, syligard 184 silicone elastomer kit, Dow Corning Corporation) mixed with a curing agent at 10:1 ratio (elastomeric base:curing agent) and incubated for 30 min. Glass coverslips were placed on the top of the negative masters containing PDMS and baked for 1h at 110° C. Pillar arrays were carefully peeled from the negative masters. The resulting radius-and length-size of individual pilus within the array was 1.79 µm and 5.8 µm, respectively. While the elasticity and the spring constant of each pilus was 2.1 MPa and 251 pN-nm, respectively, as previously reported (Patkunarajah et al., 2020; Poole et al., 2014; Servin-Vences et al., 2017). Before use for cell culture, pillar arrays were plasma cleaned with a Femto low-pressure plasma system (Deiner Electronic GmbH) and coated with EHS laminin (20 µg/mL) or Fibronectin from bovine serum (200 µg/mL).

### Electrophysiology

Whole-cell patch clamp experiments were made from DRG neurons and transiently transfected N2a^Piezo1-/-^ cells using pulled and heat-polished borosilicate glass pipettes (Harvard apparatus, 1.17 mm x 0.87 mm) with a resistance of 3-6 MΩ. The pipettes were pulled using a DMZ puller (Germany) and filled with a solution containing (in mM): 110 KCl, 10 NaCl, 1 MgCl_2_, 1 EGTA and 10 HEPES. For recordings in DRG neurons QX-314 (Alomone Labs) at 1 μM was added. The pH was adjusted to 7.3 with KOH. The extracellular solution contained (in mM): 140 NaCl, 4 KCl, 2 CaCl_2_, 1 MgCl_2_, 4 Glucose and 10 HEPES. The pH was adjusted to 7.4 with NaOH. Pipette and membrane capacitance were compensated using the auto-function of Patchmaster (HEKA, Elektornik GmbH, Germany) and series resistance was compensated to minimize voltage errors. Currents were evoked by mechanical stimuli (see below) at a holding potential of -60 mV.

Current-clamp experiments were performed to classify sensory neurons into mechano-and nociceptors. Spontaneous activity was determined by recording the membrane potential of neurons for 20 s in the absence of current injection. Cells firing action potentials in the absence of current injection were considered as a responsive cell.

For pillar arrays experiments, a single pilus was deflected using a heat-polished borosilicate glass pipette (mechanical stimulator) driven by a MM3A micromanipulator (Kleindiek Nanotechnik, Germany) as described in Poole et al., 2014(Poole et al., 2014). Only cells on the top of pili were stimulated. Pillar deflection stimuli were applied in the range of 1-1000 nm, larger deflections were discarded. For quantification and comparison analysis, the data was binned by the magnitude of the stimuli (1-10, 11-50, 51-100, 101-250, 251-500, 501-1000 nm) and calculated the mean of the current amplitudes within each bin for every cell. Bright field images (Zeiss 200 inverted microscope) were collected using a 40X objective and a CollSnapEZ camera (Photometrics, Tucson, AZ) before and after the pillar stimuli to calculate the pillar deflection. The pillar movement was calculated comparing the light intensity of the center of each pilus before and after the stimuli with a 2D-Gaussian fit (Igor Software, WaveMetrics, USA).

HSPC (Ala Scientific) experiments were carried out in excised outside-out patched pulled from transiently transfected N2a^Piezo1-/-^ cells. Recording pipettes had a final resistance of 6-8 MΩ. Positive pressure pulses were applied through the recording electrode. Pressure steps protocol consisted in ranging stimuli from 10 to 170 mmHg, in 20 mmHg steps while holding the patch potential at -60 mV. Tail currents protocol consisted of applying depolarized pre-pulses ranging from -60 to 140 mV followed by a repolarizing voltage step to -60 mV in the continuous presence of pressure. Recording solutions consisted in symmetrical ionic conditions containing (in mM): 140 NaCl, 10 HEPES, 5 EGTA adjusted to pH 7.4 with NaOH(Moroni et al., 2018). Currents were recorded at 10 KHz and filtered at 3 KHz using an EPC-10 USB amplifier (HEKA, Elektornik GmbH, Germany) and Patchmaster software. Currents and the biophysical parameters were analysed using FitMaster (HEKA, Elektornik GmbH, Germany).

### *Ex-vivo* skin nerve

Cutaneous sensory fiber recordings were performed using the *ex-vivo* skin nerve preparation. Mice were euthanized by CO_2_ inhalation for 2-4min followed by cervical dislocation. We used the recently described tibial nerve preparation to record from single-units innervating the glabrous hindpaw skin(Schwaller et al., 2021; Walcher et al., 2018a). In all preparations, the skin and nerve were dissected free and transferred to the recording chamber where muscle, bone and tendon tissues were removed from the skin to improve recording quality. The recording chamber was perfused with a 32°C synthetic interstitial fluid (SIF buffer): 123 mM NaCl, 3.5 mM KCl, 0.7mM MgSO_4_, 1.7 mM NaH_2_PO_4_, 2.0 mM CaCl_2_, 9.5 mM sodium gluconate, 5.5 mM glucose, 7.5 mM sucrose and 10 mM HEPES (pH7.4). The skin was pinned out and stretched, such that the outside of the skin could be stimulated using stimulator probes. The peripheral nerve was fed through to an adjacent chamber in mineral oil, where fine filaments were teased from the nerve and placed on a silver wire recording electrode.

The receptive fields of individual mechanoreceptors were identified by mechanically probing the surface of the skin with a blunt glass rod or blunt forceps. Analog output from a Neurolog amplifier were filtered and digitized using the Powerlab 4/30 system and Labchart 7.1 software (ADinstruments). Spike-histogram extension for Labchart 7.1 was used to sort spikes of individual units. Electrical stimuli (1Hz, square pulses of 50-500 ms) were delivered to single-unit receptive fields to measure conduction velocity and enable classification as C-fibres (velocity <1.2 ms^-1^), A-δ fibres (1.2-10 ms^-1^) or A-β fibres (>10 ms^- 1^). Mechanical stimulation of the receptive fields of neurons were performed using a piezo actuator (Physik Instrumente, P-841.60) and a double-ended Nanomotor (Kleindiek Nanotechnik, MM-NM3108) connected to a force measurement device (Kleindiek Nanotechnik, PL-FMS-LS). Calibrated force measurements were acquired simultaneously using the the Powerlab system and Labchart software during the experiment.

As different fibre types have different stimulus tuning properties, different mechanical stimuli protocols were used based on the unit type. Low threshold Aβ-fibers were stimulated with a 25 Hz vibration sitimulus with increasing amplitude over 6 steps (from ∼6-44 mN; 20 cycles per step), and a dynamic stimulus sequence with four ramp and hold waveform with varying probe deflection velocity (3 s duration; 0.075 mms^-1^, 0.15 mms^-1^, 0.45 mms^-1^ and 1.5 mms^-1^; average amplitude 100 mN). Aβ-fibre slowly-adapting mechanoreceptors (SAMs) and rapidly-adapting mechanoreceptors (RAMs) were classified by the presence or absence of firing during the static phase of a ramp and hold stimulus, respectively as previously described(Milenkovic et al., 2008; Walcher et al., 2018b). Single-units were additionally stimulated with a series of five static mechanical stimuli with ramp and hold waveforms of increasing amplitude (3 s duration; ranging from ∼10-160 mN). High threshold Aδ-and C-fibres were also stimulated using the five ramp and hold stimuli with increasing amplitudes. Spontaneous activity in C-fibres was analysed after every mechanical stimuli.

### Generation of *Piezo2*^*R2756H*^ and *Piezo2*^*R2756K*^ mice

Constitutive *knock-in* mice were generated using CRISPR-Cas9 technology by the ingenious targeting laboratory (USA). For each mutant, gRNAs (guide RNAs) and ssDNA (single-stranded DNA) donors were designed. For mutant *Piezo2*^*R2756H*^ was generated using the gRNA 5’-TGGAAGCTCTTCAAACATGATGG-3’ and the ssDNA donor 5’-TGCTGTCTCTTTCAGTATCATGGGATTGTATGCATCTGTTGTCCTTGTAATTGGGAAATTTGTTCATGAGTTCTCAGTGGGATCTCTCATTCCATCATGTTTGAAGAGCTTCCAAATGTGGACAGAATCTTGAAGTTGTGCACAGATATATTCCTCGTGAGGGAGACA-3’. Mice *Piezo2*^*R2756K*^ was generated using the gRNA 5’-TTGTTCGTGAGTTCTTCAGTGGG-3’ and the ssDNA donor 5’-ACCATCTTCATCATTTTCTCCTTGCTGTCTCTTTCAGTATCATGGGATTGTATGCATCTGTTGTCCTTGTAATTGGGAAATTTGTTAAGGAGTTCTCAGTGGGATCTCTCATTCCATCATGTTTGAAGAGCTTCCAAATGTGGA-3’. gRNAs and ssDNAs were injected into fertilized embryos (F0 mutant animals or founders). F0 embryos were transferred into pseudopregnant mice. Founders were bred with C57BL/6N mice to generate F1 mice.

### Genotyping

Ear biopsies were collected and incubated overnight at 55° C while shaking at 800 rpm in a proteinase K-lysis buffer (200 mM NaCl, 100 mM Tris pH 8.5, 5 mM EDTA, 0.2% of SDS). PCRs were performed using supernatant of the lysis preparation as DNA template (20-100 ng), 1X Taq PCR buffer, 2 mM MgCl_2_, 400 µM dNTPs, 1.25 U Taq-polymerase (Thermofisher Scientific) and 0.5 µM of primers. A 499 bp fragment of *Piezo2* locus was amplified using the forward 5’-GAAAGAGCTACTTTGAAAGGAGTATGTGC-3’ and reverse 5’-CCTGTCAGAAGAGAAATGGTTGCC-3’ primers. Inserted point mutations generated new restrictions sites that allow to identify wild type, heterozygous and homozygous animals from each knock-in mice. PCR products were incubated overnight with BspI and MseI restriction endonucleases (NEB Inc.) for *Piezo2*^*R2756H*^ and *Piezo2*^*R2756K*^ mouse lines, respectively. Amplified and digested DNA fragments were observed by gel electrophoresis.

### RNAscope

Lumbar DRGs were collected from adult animals and incubated for 40 min in Zamboni’s fixative media (2% of para-formaldehyde + picric-acid), washed with PBS and incubated in 30% sucrose (in PBS) overnight at 4°C. DRGs were embedded in OCT Tissue Tek (Sakura, Alphen aan den Rijn). 10 µm-thick cryosections were stored at -80°C until used for experiments. In situ hybridization was carried out according to the manufacturer’s instructions (RNAscope™ Multiplex Fluorescent V2 assay, ADC, Kit #323110, *Piezo2* probe #4001191). LSM700 Carls Zeiss confocal microscope was use to acquired images at 20X and numerical aperture 0.5. Fluorescence intensity was analysed using ImageJ.

### Brush and von Frey experiments

For both methods, testing was carried out during the light phase. Males and females were included in the experiments. Mice (7-12 weeks old) were placed in plastic cages with a metal grid floor bottom that allowed access to hindpaw stimulation. Animals were habituated for two-consecutives days for 20 min before testing. Before starting the behavioral test, animals were placed in the cages for at least 20 min (accommodation) and experimentation started when animals stopped cage exploration. Each animal was tested at least twice on different days and average values from those measures were plotted in each animal.

Responses to gentle touch was measured by stroking the surface of the hindpaw. Animals were stimulated five times with at least 2 min between each stimulation. The percentage was calculated according to the number of withdrawals out of the five stimulations.

Calibrated von Frey (Semmes-Weinstein) filaments (Aesthesio®, USA) were used to test the 50% paw withdrawal threshold (50% PWT) in mice. The “up-down method” was used to calculate the 50% PWT(Chaplan et al., 1994; Christensen et al., 2020; Dixon, 1980). Mid-plantar right and left hindpaw were stimulated for approximately 1-3 s with von Frey filaments in the range of 0.008, 0.02, 0.04, 0.07, 0.16, 0.4, 0.6, 1.0, 1.4 and 2 g. Von Frey filaments were presented perpendicular to hind paws at intervals of 3-5 min. A positive response was considered when the paw was withdrawn.

## Statistical analysis

All data analyses were performed using GraphPad Prism and all data sets were tested for normality. Parametric data sets were compared using a two-tailed, Student’s *t*-test. Nonparametric data sets were compared using a Mann-Whitney test. To compare more than two groups, One-way ANOVA was used. Categorical data were compared using χ^2^ tests.

## Acknowledgments

We thank Maria Braunschweig, Franziska Bartelt, and Kathleen Barda for technical assistance.

## Author Contributions

Conceptualization OSC, VB and GRL; Investigation OSC, SC, JK, FS and VB; Project administration VB; Funding acquisition GRL Supervision GRL; Writing original draft OSCand GRL, Writing review and editing OSC, VB and GRL.

## Data and materials availability

Raw data from this manuscript is available upon request.

