## Supplementary Material and Figures for "Piezo2 voltage-block regulates mechanical pain sensitivity"

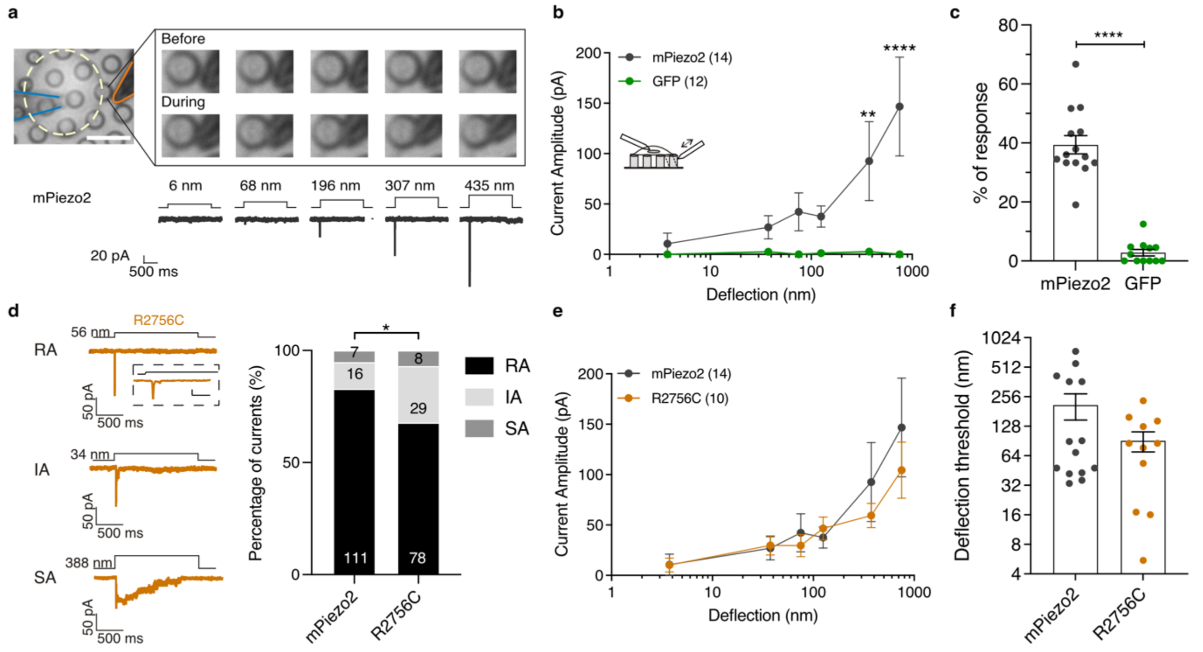


**Supplementary Fig 1. R2756C variant displayed less RA deflection-gated currents compared to control in $N2a^{Piezo1-/-}$**

**a**, Bright-field picture of a $N2a^{Piezo1-/-}$ cell overexpressing mPiezo2 and cultured on pillar arrays. The insert shows an amplification of the stimulated pilus and the sequential stimuli applied with their corresponding deflection-gated inward current (grey traces). Traces correspond to RA currents. Note that the larger the deflection, the larger the MA current. **b**, Stimulus-response plot of the deflection sensitive currents from $N2a^{Piezo1-/-}$ overexpressing mPiezo2 (gray) and GFP (green) (Two-way ANOVA, Sidak's multiple comparison test; **P=0.002, ****P<0.0001). **c,** Percentage of response to deflection stimuli (39.39 ± 3.08 and 2.8 ± 1.09 % for mPiezo2- and GFP-transfected $N2a^{Piezo1-/-}$ cells, respectively). The total amount of stimuli was considered as 100%. Each dot represents the percentage of individual cells (Student's t test; ****P<0.0001). **d**, *Left*, Representative traces of the three types of deflection-gated currents from $N2a^{Piezo1-/-}$ cells overexpressing R2765C mutant. Rapidly adapting (RA), intermediate adapting (IA) and slowly adapting (SA) currents are shown. *Right*, Proportion of RA currents is decreased in R2756C variant. Numbers in the histograms show the number of currents recorded ($\chi^2$ test, *P=0.01). **e**, Deflection-current amplitude relationship of mPiezo2 and R2756C mutant. **f,** Histogram showing that mPiezo2 and R2756C variant showed similar deflection thresholds. Data was plotted as mean ± s.e.m.

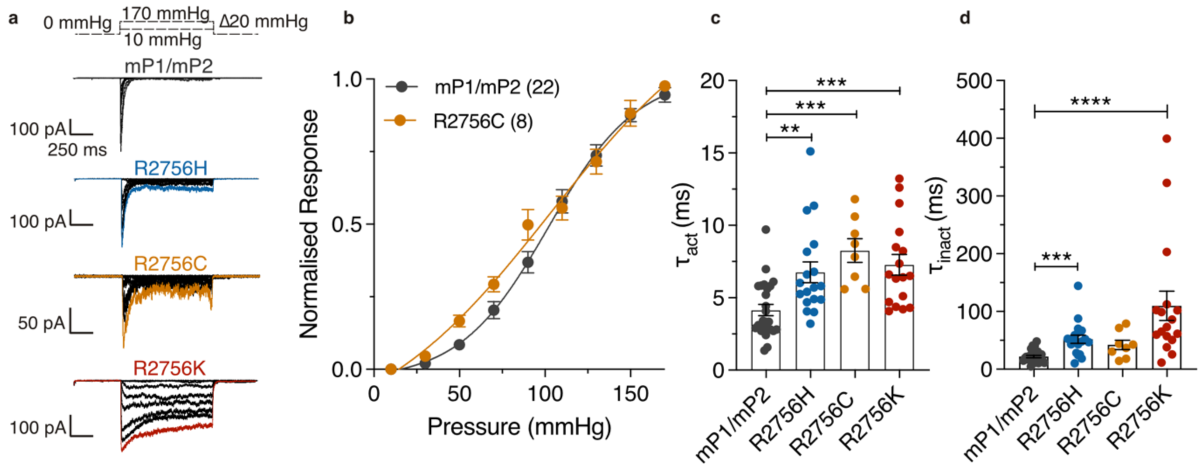


**Supplementary Fig 2. Stretch-sensitive currents from mP1/mP2 variants displayed altered activation and inactivation kinetics.**

**a**, Representative recordings of stretch-sensitive currents of outside-out patches clamped at -60 mV from N2a$^{Piezo1-/-}$ cells overexpressing the chimeric channel mP1/mP2 and mutants. The dashed lines above show the pressure protocol applied. **b**, Stretch-response curves and mechanical sensitivity of mP1/mP2 and R2756C chimeric channels. The peak currents were normalised according to the maximum amplitude current recorded. **c-d**, Time constant of activation ($\tau_{act}$, **c**), and inactivation ($\tau_{inact}$, **d**) change in mutants of the chimeric channels. The values correspond to the currents recorded at 130 mmHg pulse (Kruskal-Wallis test, *P=0.01; ***P<0.001, ****P<0.0001). In all cases, values were plotted as mean ± s.e.m. Each dot represents the value of single cell measurements.

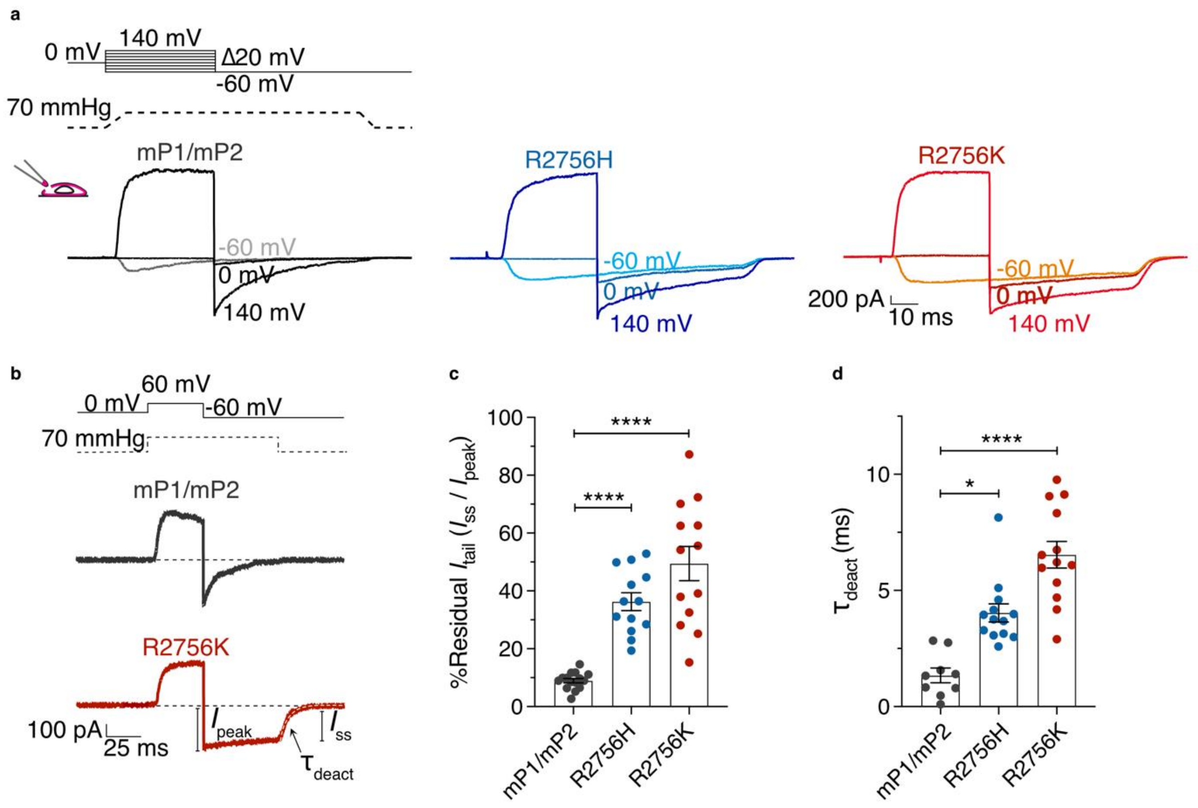


**Supplementary Fig. 3. Channel availability and deactivation change in the stretch-sensitive mutants of mP1/mP2 channels.**

**a**, Example traces of the tail current protocol performed in N2a$^{Piezo1-/-}$ cells overexpressing the chimeric variants in presence of 70 mmHg of pressure. Note that the tail currents in the mutants are larger at -60 and 0 mV compared to the control, indicating increased channel availability. **b**, Representative traces from outside-out patches clamped to a voltage step of +60 mV followed by a step of -60 mV in the presence of pressure stimuli of 70 mmHg. Peak ($I_{peak}$) and steady-state ($I_{ss}$) currents are indicated. **c**, Ratio of $I_{peak}$ to $I_{ss}$ from the instantaneous tail currents ($I_{tail}$) at -60 mV (Dunnett test; ****P<0.0001). **d**, Kinetics of the inactivation-deactivation state ($\tau_{deact}$) from mP1/mP2 variants (Kruskal-Wallis test; *P=0.01, ****P<0.0001). Data was plotted as mean ± s.e.m.

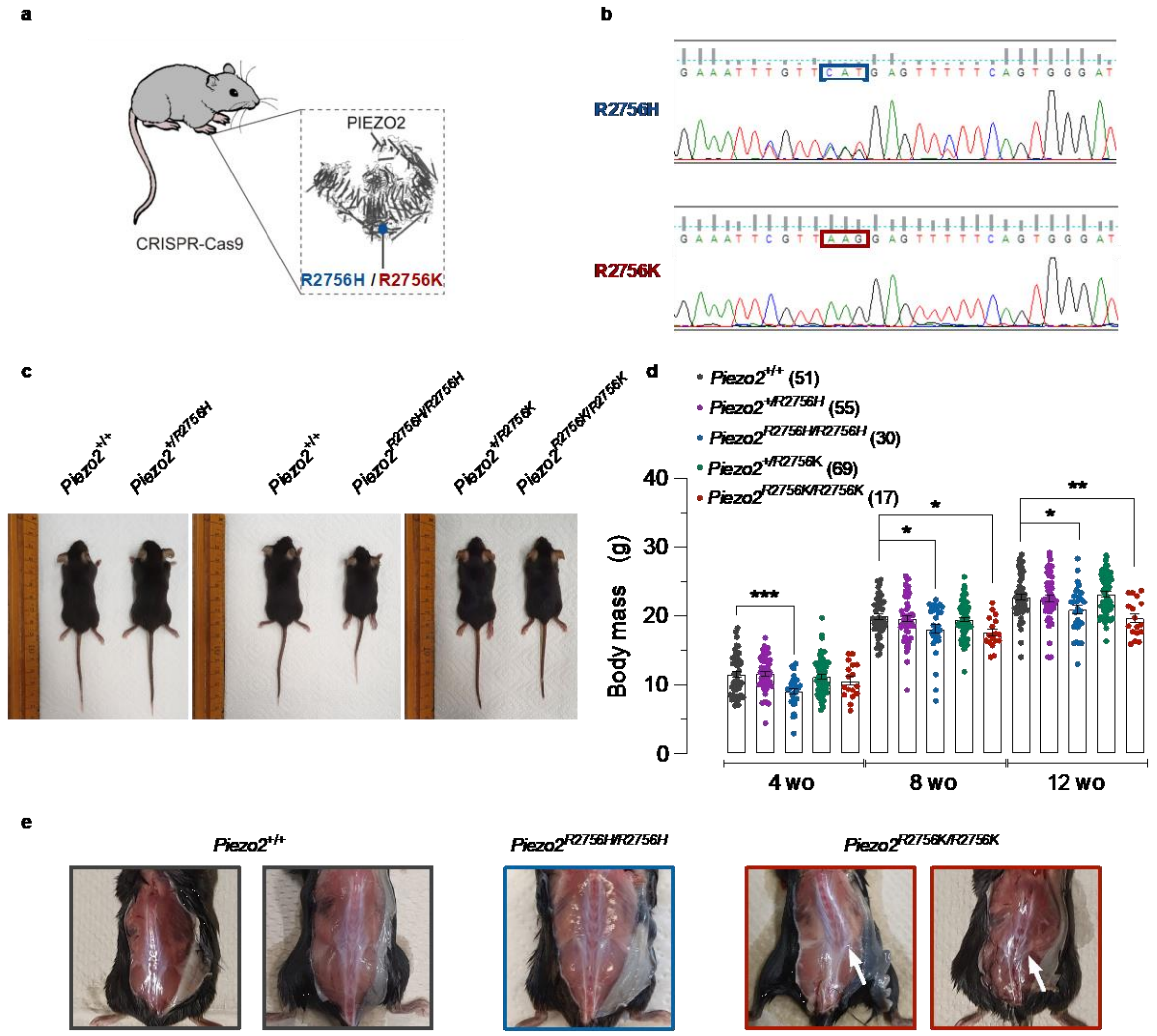


**Supplementary Fig. 4. *Piezo2 knock-in* mice showed reduced weight and scoliosis**

**a**, Cartoon indicating the global insertion of mutations R2756H and R2756K in Piezo2. **b**, Sequencing results from founder mice carrying the mutations R2756H (*above*) and R2756K (*below*) in Piezo2. The substituted nucleotides are highlighted inside the squares. Note that in *Piezo2* wild type animals Arginine is encoded by CGT codon (*not shown*). **c**, Photos of *knock-in* mice at 5 weeks old. Note that *Piezo2*$^{R2756H/R2756H}$ mice are smaller than controls. **d**, Bar plot showing that *Piezo2*$^{R2756H/R2756H}$ mice are smaller at week four after birth and that both *Piezo2*$^{R2756H/R2756H}$ and *Piezo2*$^{R2756K/R2756K}$ showed reduced weight at weeks 8 and 12 . Each dot represents an animal (mean ± s.e.m.; One-Way ANOVA test; *P<0.05, **P<0.01, ***P=0.0003). **e**, Pictures of *Piezo2*$^{+/+}$, *Piezo2*$^{R2756H/R2756H}$ and *Piezo2*$^{R2756K/R2756K}$ mice. 9 out of 18 examined *Piezo2*$^{R2756K/R2756K}$ animals showed scoliosis.

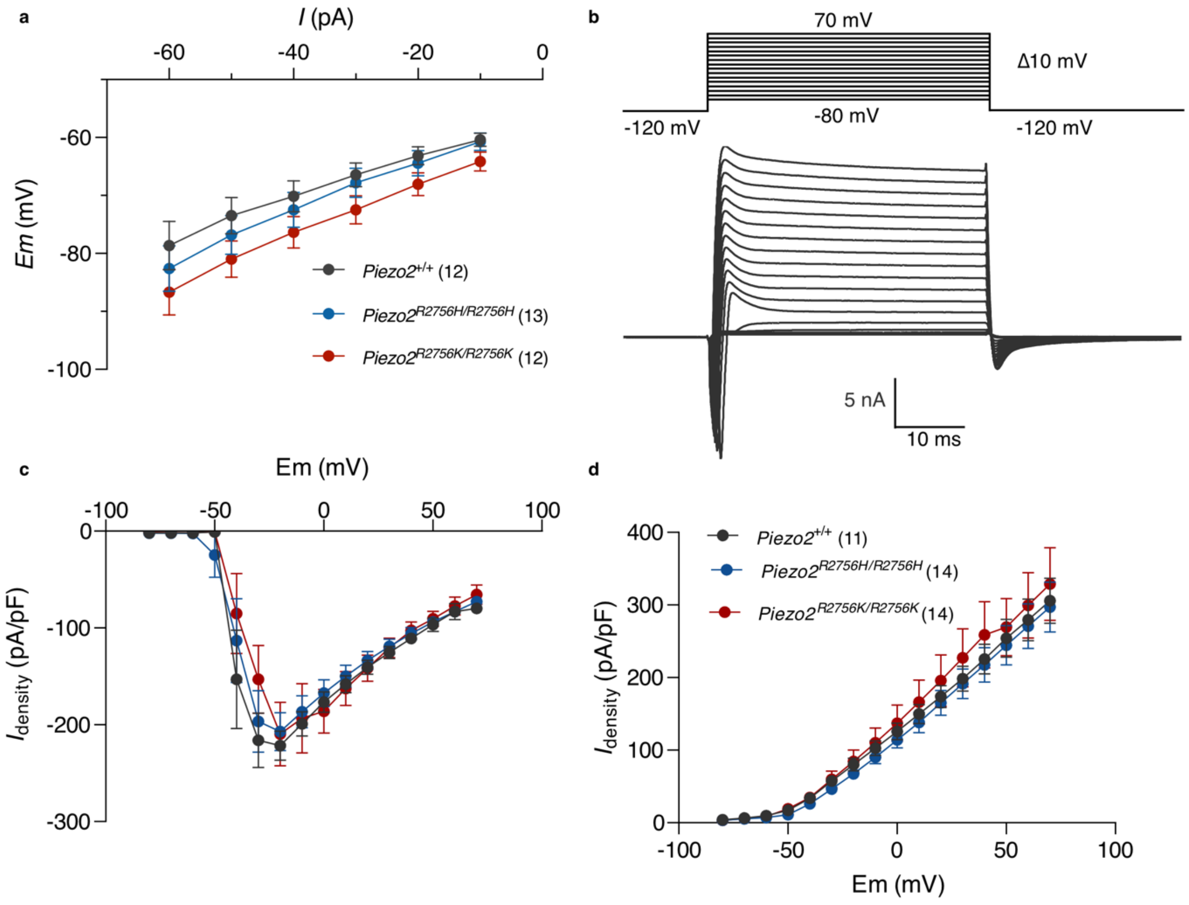


**Supplementary Fig. 5. Macroscopic inward and outward currents do not change in mechanoreceptors from *Piezo2* mutants**

**a,** Current (*I*) -voltage (Em) plot showing that input resistance did not change in mechanoreceptors from *Piezo2 knock-in* mice. **b**, Representative traces from inward and outward currents recorded from a wild type mechanoreceptor. **c, d**, Voltage (Em) - Current density ($I_{density}$) relationships showing that inward currents (**c**) and outward currents (**d**) from mechanoreceptors were similar in *Piezo2*$^{+/+}$ and mutants. Data was plotted as mean ± s.e.m.

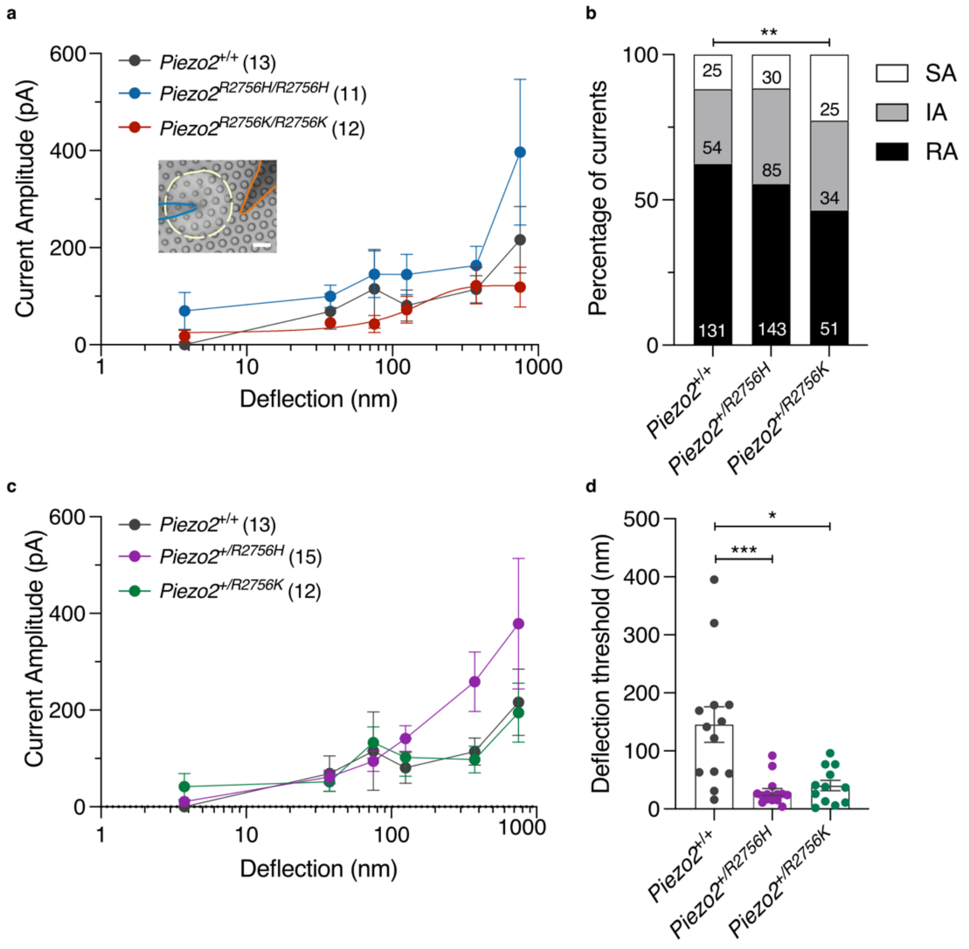


**Supplementary Fig 6. Heterozygous conditions of *Piezo2 knock-in* mice showed more sensitive deflection-gated currents in mechanoreceptors.**

**a**, Deflection-current amplitude relationship of mechanoreceptors from *knock-in* and control mice. Note that neurons from *Piezo2*$^{R2756K/R2756K}$ reached current saturation, indicating increased sensitivity to mechanical stimuli. **b**, Histograms showing that *Piezo2*$^{+/R2756K}$ mechanoreceptors evoked less RA currents compared to wild type cells ($\chi^2$ test, **P=0.009). Numbers indicate the total of currents recorded. **c**, Deflection-current amplitude plot showing no differences between wild type mechanoreceptors and neurons from both heterozygous mice. **d**, Deflection thresholds were lower in mechanoreceptors from both heterozygous mice compared to wild type cells. (Kruskal-Wallis test, *P=0.01, ***P=0.0004) Data was plotted as mean ± s.e.m.

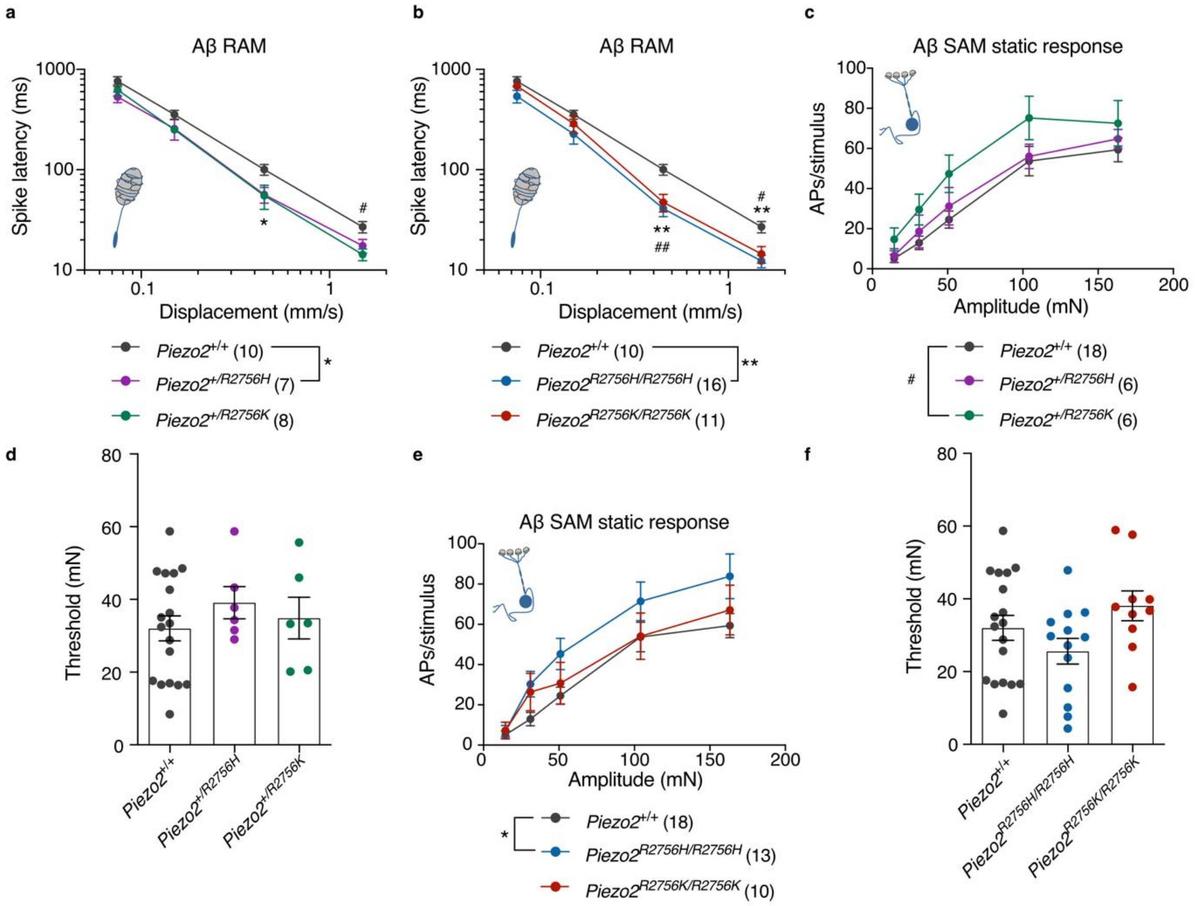


**Supplementary Fig. 7. Mechanical thresholds from Aβ SAM fibres were not affected in *Piezo2 knock-in* mice.**

**a,b**, Rapidly adapting afferents from *Piezo2 knock-in* mice displayed decreased AP firing latencies at velocity stimuli indicating increased sensitivity to mechanical stimuli (Two-way ANOVA with Sídák post hoc analysis; **a**, *P<0.05, $^{\#}$P=0.01; **b**, **P<0.01, $^{\#}$P=0.03, $^{\#\#}$P=0.009). **c,d**, Slowly adapting fibres showed a slightly AP firing increase in *Piezo2*$^{+/R2756K}$ mutants (Two-way ANOVA with Sídák post hoc analysis, $^{\#}$P=0.01), but mechanical thresholds were not impaired compared to wild type. **e,f**, SAM afferents from *Piezo2*$^{R2756H/R2756H}$ displayed higher AP firing compared to wild type fibres during sustained mechanical stimuli (Two-way ANOVA with Sídák post hoc analysis, *P=0.01), but no differences in mechanical thresholds were observed.

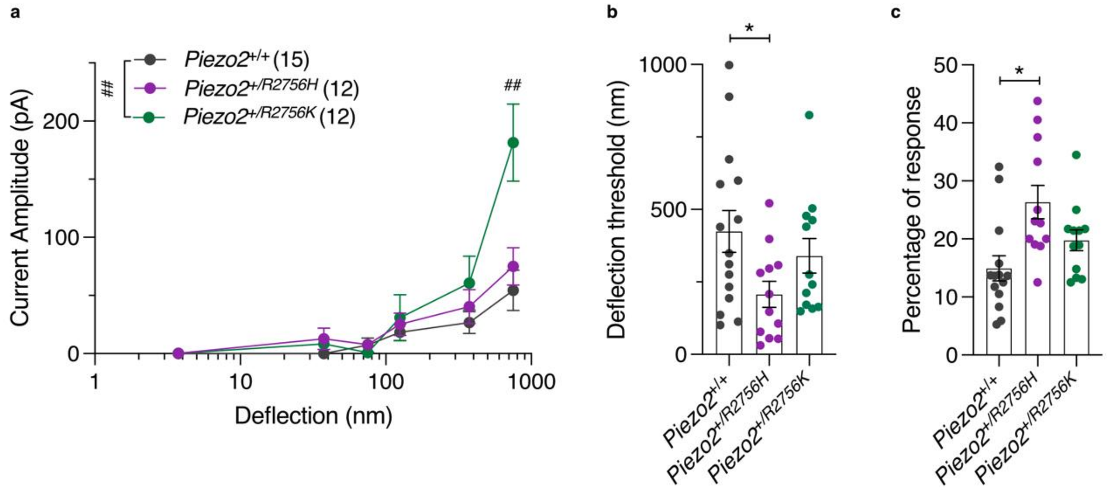


**Supplementary Fig. 8. *Piezo2$^{+/R2756H}$* nociceptors displayed more sensitive deflection-gated currents than controls.**

**a**, Deflection-current amplitude relationships of nociceptors from heterozygous *Piezo2 knock-in* and control mice (Mann Whitney test $^{\#\#}$P=0.003; additionally an ordinary Two-way ANOVA indicated differences between *Piezo2$^{+/+}$* and *Piezo2$^{+/R2756K}$*, $^{\#\#}$P=0.003). **b**, Plots showing that deflection thresholds were lower in *Piezo2$^{+/R2756H}$* neurons compared to wild type (One-Way ANOVA, *P=0.03). **c**, Dot plot showing that *Piezo2$^{+/R2756H}$* cells showed enhanced responsiveness to deflection stimuli compared to controls (One-Way ANOVA, *P=0.03). Data was plotted as mean ± s.e.m.

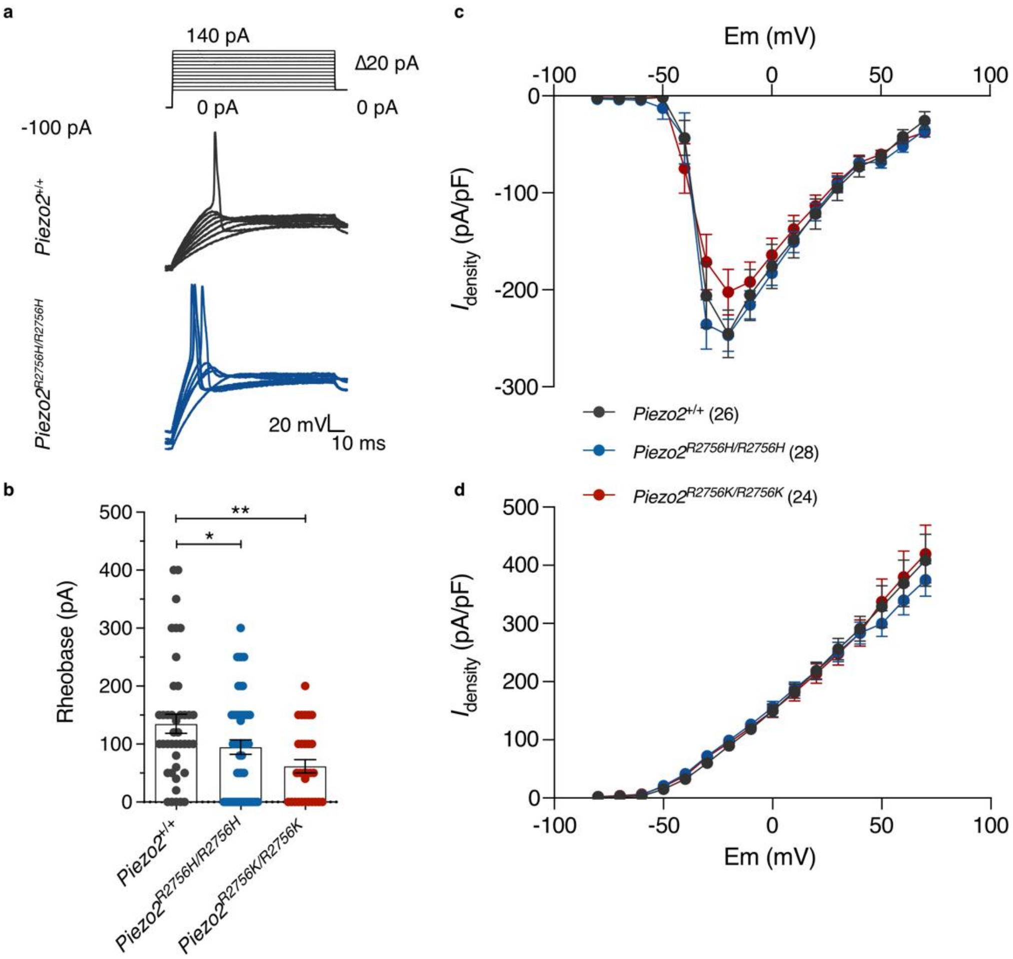


**Supplementary Fig. 9. Nociceptors from *Piezo2 knock-in* mice displayed hyperexcitability**

**a**, Representative traces from APs evoked at different current step injections in nociceptors from *Piezo2*$^{+/+}$ and *Piezo2*$^{R2756H/R2756H}$. Note that *Piezo2*$^{R2756H/R2756H}$ cells showed lower thresholds. **b**, Dot plots showing that *Piezo2 knock-in* neurons exhibited lower AP threshold firings (Rheobase) (One-way ANOVA, *P=0.03; **P=0.001). **c, d**, Voltage (Em)-Current density ($I_{density}$) relationships showing that inward currents (**c**) and outward currents (**d**) from nociceptors were similar in *Piezo2*$^{+/+}$ and mutants. Data was plotted as mean ± s.e.m.

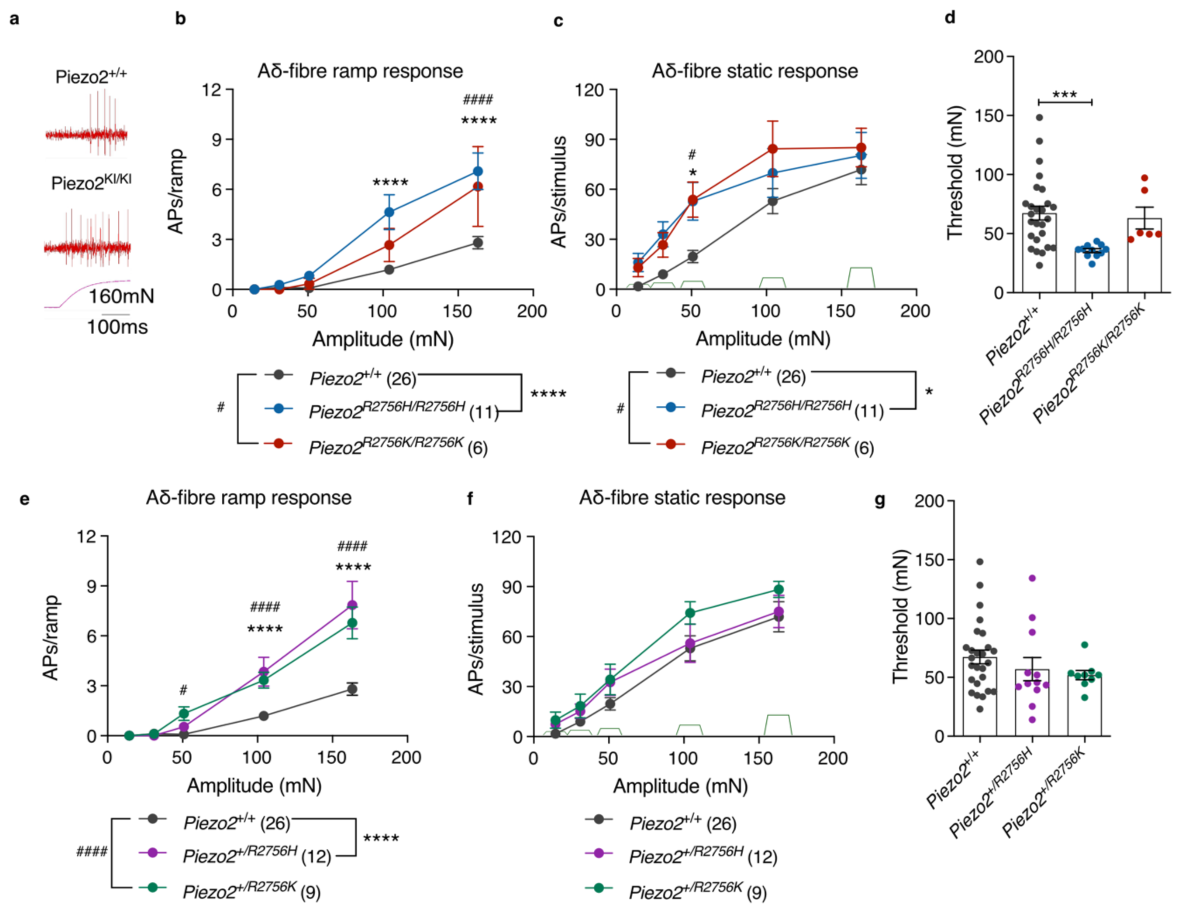


**Supplementary Fig. 10. Aδ-nociceptor firing is enhanced in *Piezo2 knock-in* mice**
**a**, Example traces of responses to ramp phases in Aδ-nociceptor from *Piezo2*$^{+/+}$ and *Piezo2 knock-in* animals during a 160 mN stimulus. **b,e**, Aδ-fibre firing activity during ramp phase (Two-way ANOVA with Sídák post hoc analysis, *P<0.05, ****P<0.0001, $^{\#\#\#\#}$P<0.0001). Note that all mutants showed enhanced firing compared to controls. **c,f**, Static phase responses from Aδ-fibres in *Piezo2*$^{+/+}$ and mutants (Two-way ANOVA with Sídák post hoc analysis, *P<0.05, $^{\#}$P<0.05). Only homozygous displayed higher firing activity at the 31 mN force. **d,g**, Dot plot showing that Aδ-nociceptors from *Piezo2*$^{R2756H/R2756H}$ are more sensitive to mechanical stimuli compared to controls (Kruskal-Wallis, ***P<0.001). Data are presented as mean ± s.e.m.

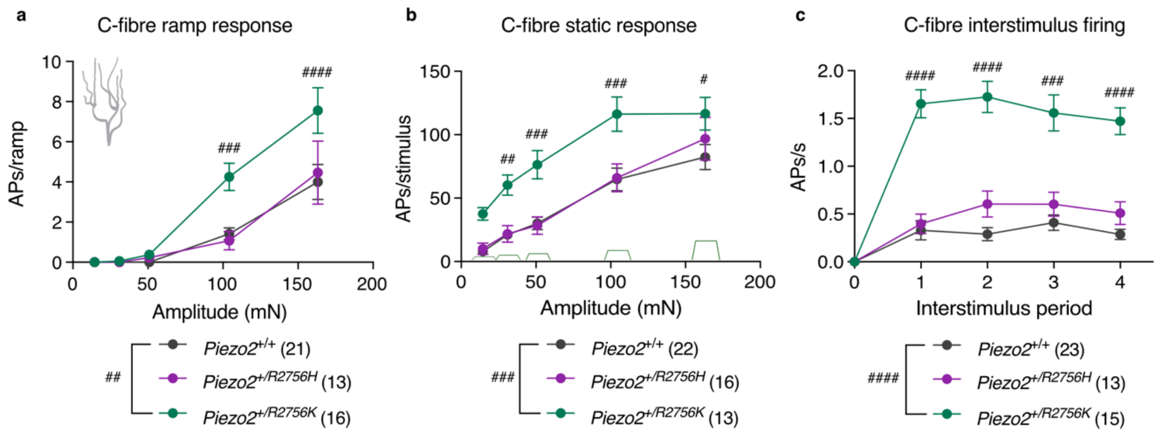


**Supplementary Fig. 11. *Piezo2$^{+/R2756K}$* nociceptors exhibited after discharge firing activity**

**a-c**, AP firing activity of C-fibres during ramp phase (**a**, Two-way ANOVA with Sídák post hoc analysis, $^{\#\#}$P=0.001, $^{\#\#\#}$P=0.0004, $^{\#\#\#\#}$P<0.0001), static phase (**b**, Two-way ANOVA with Sídák post hoc analysis, $^{\#}$P=0.02, $^{\#\#}$P=0.006, $^{\#\#\#}$P<0.001) and interstimulus firing (**c**, Two-way ANOVA with Sídák post hoc analysis, $^{\#\#\#}$P=0.0001, $^{\#\#\#\#}$P<0.0001). Data are presented as mean ± s.e.m.

| | mPiezo2 | R2756H | R2756C | R2756K |
|---|---|---|---|---|
| **Cells (no. of currents)** | 14 (134) | 11 (130) | 10 (115) | 11 (172) |
| **Latency (ms)** | **1.14 ± 0.62** | **2.47 ± 0.37**** | **2.54 ± 0.37**** | **2.34 ± 0.27** |
| **RA** | 1.2 ± 0.83 | 2.44 ± 0.55 | 2.07 ± 0.35 | 2.32 ± 0.35 |
| **IA** | 1.35 ± 0.28 | 2.47 ± 0.54 | 3.34 ± 0.65 | 2.61 ± 0.76 |
| **SA** | 1.29 ± 0.41 | 3.2 ± 0.65 | 7.76 ± 3.9 | 2.67 ± 0.81 |
| **$\tau_{act}$ (ms)** | **0.65 ± 0.04** | **0.86 ± 0.05**** | **0.68 ± 0.04** | **0.74 ± 0.06** |
| **RA** | 0.60 ± 0.05 | 0.72 ± 0.05 | 0.58 ± 0.03 | 0.62 ± 0.04 |
| **IA** | 1.06 ± 0.17 | 1.22 ± 0.13 | 0.97 ± 0.19 | 1.11 ± 0.27 |
| **SA** | 1.14 ± 0.23 | 0.81 ± 0.18 | 0.9 ± 0.25 | 0.96 ± 0.16 |
| **$\tau_{inact}$ (ms)** | **14.02 ± 5.64** | **21.08 ± 5.86***** | **18.49 ± 6.35** | **16.51 ± 3.27** |
| **RA** | 0.83 ± 0.08 | 1.76 ± 0.1* | 1.04 ± 0.12 | 0.99 ± 0.1 |
| **IA** | 17.62 ± 3.28 | 16.74 ± 2.01 | 15.25 ± 1.75 | 17.99 ± 3.07 |
| **SA** | 212.2 ± 75.35 | 198.2 ± 50.76 | 170.7 ± 56.35 | 114.4 ± 17.02 |

**Extended Data Table 1.**

**Biophysical properties of N2a$^{Piezo1-/-}$ cells overexpressing mPiezo2 mutants**

Number of cells and currents analysed from cells overexpressing pathogenic mutations of Piezo2. All data come from, at least, three different transfections. Mechanical latency, $\tau_{act}$ and $\tau_{inact}$ are shown for all cells including the values or RA, IA and SA groups. (Kruskal-Wallis test, *P=0.04, **P<0.01,***P<0.001). Values are shown as mean ± s.e.m.

| | Piezo2$^{+/+}$ | Piezo2$^{+/R2756H}$ | Piezo2$^{R2756H/R2756H}$ | Piezo2$^{+/R2756K}$ | Piezo2$^{R2756K/R2756K}$ |
|---|---|---|---|---|---|
| **Animals (m,f)** | 6 (5,1) | 7 (6,1) | 6 (4,2) | 4 (3,1) | 5 (3,2) |
| **Cells (no. of currents)** | 13 (210) | 15 (258) | 11 (145) | 12 (110) | 12 (123) |
| **Cell size (µm)** | 37.96 ± 0.98 | 41.33 ± 0.69 | 37.3 ± 1.05 | 39.44 ± 1.96 | 36.87 ± 0.68 |
| **HP (ms)** | 0.66 ± 0.05 | 0.65 ± 0.12 | 0.6 ± 0.11 | 0.7 ± 0.08 | 0.61 ± 0.08 |
| **$Em_{rep}$ (mV)** | -66.72 ± 1.04 | -66.78 ± 1.64 | -65.16 ± 2.33 | -64.89 ± 1.08 | -63.13 ± 1.31 |
| **Latency (ms)** | **3.19 ± 0.42** | **2.77 ± 0.34** | **4.98 ± 0.64** | **5.23 ± 0.78****** | **3.86 ± 0.45****** |
| **RA** | 3.98 ± 0.9 | 1.73 ± 0.3 | 4.79 ± 0.87 | 4.73 ± 1.07** | 3.21 ±0.53* |
| **IA** | 3.10 ± 0.87 | 4.49 ± 0.86 | 4.64 ± 1.18 | 5.05 ± 1.39 | 5.57 ± 1.25** |
| **SA** | 2.22 ± 0.81 | 2.01 ± 0.47 | 5.92 ± 1.81 | 5.52 ± 1.62 | 4.18 ± 0.91 |
| **$\tau_{act}$ (ms)** | **0.87 ± 0.04** | **1.2 ± 0.06**** | **1.43 ± 0.09****** | **1.52 ± 0.09****** | **1.33 ± 0.11***** |
| **RA** | 0.71 ± 0.04 | 0.84 ± 0.05 | 1.27 ± 0.13**** | 1.25 ± 0.11**** | 0.97 ± 0.07** |
| **IA** | 1.07 ± 0.09 | 1.72 ± 0.15** | 1.57 ± 0.19 | 1.65 ± 0.15** | 1.96 ± 0.35* |
| **SA** | 1.26 ± 0.12 | 1.46 ± 0.21 | 1.62 ± 0.23 | 1.85 ± 0.26 | 1.96 ± 0.32 |
| **$\tau_{inact}$ (ms)** | **23.86 ± 3.83** | **31.43 ± 6.05** | **42.37 ± 8.96**** | **61.89 ± 14.67**** | **32.2 ± 8.98** |
| **RA** | 1.79 ± 0.1 | 1.82 ± 0.11 | 2.43 ± 0.16** | 1.95 ± 0.14 | 1.94 ± 0.14 |
| **IA** | 18.8 ± 1.57 | 16.16 ± 1.32 | 14.02 ± 1.36* | 17.35 ± 2.16 | 13.48 ± 1.75 |
| **SA** | 149.5 ± 16.8 | 215.9 ± 37.45 | 184.4 ± 34.45 | 246.4 ± 50.27 | 181.4 ± 45.45 |

**Extended Data Table 2.**

**Biophysical properties of deflection gated currents in mechanoreceptors from *Piezo2$^{R2756H}$* and *Piezo2$^{R2756K}$* mice**

Sex and number of animals (m, male, f, female) used in the study. Cells and currents analysed in mechanoreceptors for each genotype are shown. Cell size and half peak (HP) of APs were used to classified DRG neurons into mechanoreceptors. Mechanical latency, , $\tau_{act}$ and $\tau_{inact}$ are shown for all genotypes including the values or RA, IA and SA groups. (Kruskal-Wallis test, *P<0.05, **P<0.01, ***P<0.001, ****P<0.0001). Values are shown as mean ± s.e.m.

| | Piezo2$^{+/+}$ | Piezo2$^{+/R2756H}$ | Piezo2$^{R2756H/R2756H}$ | Piezo2$^{+/R2756K}$ | Piezo2$^{R2756K/R2756K}$ |
|---|---|---|---|---|---|
| **Animals (m,f)** | 5 (4,1) | 4 (3,1) | 4 (2,2) | 3 (2,1) | 4 (3,1) |
| **Cells (no. of currents)** | 15 (77) | 12 (62) | 13 (97) | 12 (54) | 14 (113) |
| **Cell size (μm)** | 26.67 ± 1.13 | 23.61 ± 0.66 | 26.04 ± 1.32 | 24.84 ± 0.72 | 24.45 ± 0.86 |
| **HP (ms)** | 2.21 ± 0.27 | 2.54 ± 0.36 | 2.59 ± 0.31 | 1.75 ± 0.17 | 2.18 ± 0.25 |
| **$Em_{rep}$ (mV)** | -54.6 ± 2.6 | -52.5 ± 2.7 | -52.9 ± 3.1 | -54.7 ± 2.4 | -49.7 ± 3.2 |
| **Latency (ms)** | **1.44 ± 0.16** | **2.24 ± 0.45** | **2.41 ± 0.43** | **2.96 ± 0.55*** ** | **2.68 ± 0.34** ** |
| **RA** | 1.25 ± 0.15 | 1.85 ± 0.53* | 2.31 ± 0.59 | 2.29 ± 0.49** | 2.28 ± 0.44** |
| **IA** | 1.81 ± 0.53 | 2.2 ± 0.56 | 2.78 ± 0.76 | 4.41 ± 1.14 | 2.73 ± 0.53 |
| **SA** | 2.29 ± 0.77 | 2.95 ± 1.28 | 2.25 ± 0.66 | 3.7 ± 1.63 | 3.96 ± 0.92 |
| **$\tau_{act}$ (ms)** | **0.75 ± 0.07** | **0.83 ± 0.05** | **1.00 ± 0.11** | **1.02 ± 0.09** ** | **1.24 ± 0.13*** |
| **RA** | 0.6 ± 0.05 | 0.67 ± 0.8 | 0.63 ± 0.05 | 0.77 ± 0.09 | 0.73 ± 0.08 |
| **IA** | 0.89 ± 0.17 | 0.85 ± 0.1 | 1.97 ± 0.4 | 1.39 ± 0.25 | 1.37 ± 0.19 |
| **SA** | 1.86 ± 0.34 | 1.08 ± 0.08 | 1.18 ± 0.26 | 1.1 ± 0.19 | 2.61 ± 0.55 |
| **$\tau_{inact}$ (ms)** | **17.32 ± 5.02** | **69.29 ± 19.78****** | **22.15 ± 5.78** | **62.41 ± 16.36*** ** | **45.57 ± 16.44** ** |
| **RA** | 0.9 ± 0.1 | 1.58 ± 0.23 | 1.32 ± 0.17 | 1.6 ± 0.26 | 1.47 ± 0.14 |
| **IA** | 16.92 ± 3.29 | 15.59 ± 2.84 | 22.71 ± 2.52 | 21.91 ± 3.56 | 16.56 ± 3.26 |
| **SA** | 136.9 ± 22.05 | 235.3 ± 55.26 | 142.2 ± 33.37 | 230.8 ± 41.94 | 257.4 ± 85.34 |

### Extended Data Table 3.

**Biophysical properties of deflection gated currents in nociceptors from *Piezo2$^{R2756H/K}$* mice**

Sex and number of animals (m, male, f, female) used in the study. Cells and currents analysed in nociceptors for each genotype are shown. Cell size and half peak (HP) of APs were used to classified DRG neurons into nociceptors. Mechanical latency, $\tau_{act}$ and $\tau_{inact}$ are shown for all genotypes including the values or RA, IA and SA groups. (Kruskal-Wallis test, *P<0.05, **P<0.01,***P<0.001, ****<0.0001). Values are shown as mean ± s.e.m.
